# Automated and accurate estimation of gene family abundance from shotgun metagenomes

**DOI:** 10.1101/022335

**Authors:** Stephen Nayfach, Patrick H. Bradley, Stacia K. Wyman, Timothy J. Laurent, Alex Williams, Jonathan A. Eisen, Katherine S. Pollard, Thomas J. Sharpton

**Affiliations:** Gladstone Institute of Cardiovascular Disease, San Francisco, CA, USA; Department of Evolution and Ecology, Department of Medical Microbiology and Immunology, UC Davis Genome Center, University of California, Davis; Department of Epidemiology & Biostatistics and Institute for Human Genetics, University of California, San Francisco, CA, USA; Department of Microbiology, Department of Statistics, Oregon State University, Corvallis, OR, USA

## Abstract

Shotgun metagenomic DNA sequencing is a widely applicable tool for characterizing the functions that are encoded by microbial communities. Several bioinformatic tools can be used to functionally annotate metagenomes, allowing researchers to draw inferences about the functional potential of the community and to identify putative functional biomarkers. However, little is known about how decisions made during annotation affect the reliability of the results. Here, we use statistical simulations to rigorously assess how to optimize annotation accuracy and speed, given parameters of the input data like read length and library size. We identify best practices in metagenome annotation and use them to guide the development of the Shotgun Metagenome Annotation Pipeline (ShotMAP). ShotMAP is an analytically flexible, end-to-end annotation pipeline that can be implemented either on a local computer or a cloud compute cluster. We use ShotMAP to assess how different annotation databases impact the interpretation of how marine metagenome and metatranscriptome functional capacity changes across seasons. We also apply ShotMAP to data obtained from a clinical microbiome investigation of inflammatory bowel disease. This analysis finds that gut microbiota collected from Crohn’s disease patients are functionally distinct from gut microbiota collected from either ulcerative colitis patients or healthy controls, with differential abundance of metabolic pathways related to host-microbiome interactions that may serve as putative biomarkers of disease.

**Author Summary:** Microbial communities perform a wide variety of functions, from marine photosynthesis to aiding digestion in the human gut. Shotgun “metagenomic” sequencing can be used to sample millions of short DNA sequences from such communities directly, without needing to first culture its constituents in the laboratory. Using these data, researchers can survey which functions are encoded by mapping these short sequences to known protein families and pathways. Several tools for this annotation already exist. But, annotation is a multi-step process that includes identification of genes in a metagenome and determination of the type of protein each gene encodes. We currently know little about how different choices of parameters during annotation influences the final results. In this work, we systematically test how several key decisions affect the accuracy and speed of annotation, and based on these results, develop new software for annotation, which we named ShotMAP. We then use ShotMAP to functionally characterize marine communities and gut communities in a clinical cohort of inflammatory bowel disease. We find several functions are differentially represented in the gut microbiome of Crohn’s disease patients, which could be candidates for biomarkers and could also offer insight into the pathophysiology of Crohn’s. ShotMAP is freely available (https://github.com/sharpton/shotmap).

## Introduction

Sequencing DNA collected from microbial communities has been critical to the study of uncultured microorganisms. High-throughput amplicon sequencing of taxonomically informative loci (e.g., SSU-rRNA genes) has shed light on the tremendous diversity and distribution of microbial communities in nature [1] and revealed patterns and processes related to the assembly [2], diversification [3], and scaling [4] of these communities. Shotgun sequencing of total DNA from microbial communities (*i.e.,* metagenomics) is gaining popularity, as it provides insight into the genomic composition of microbes as they exist in nature and enables inference of the community’s biological functional potential [5]. By annotating metagenomic sequences with the gene families from which they derive, the community’s biological functional potential can be profiled. Comparing these profiles to other metagenomes or to environmental covariates enables (i) quantification of how functions vary across samples, (ii) identification of functions that correlate with ecological parameters, and (iii) discovery of functions that stratify communities (*i.e.,* biomarkers).

Several methods of functionally annotating metagenomes have been developed. These include stand-alone software such as MEGAN [6], HUMAnN [7], RAMMCAP [8], SmashCommunity [9], and MOCAT [10], as well as cloud-based tools like CloVR [11], and web portals like MG-RAST [12], MicrobesOnline [13], and the IMG/M annotation server [14][S1 Table]. Generally, these methods operate by comparing metagenomic sequence reads to a reference database of functionally annotated protein families and use homology inference to annotate each read [5]. Despite the wide use of this general strategy, surprisingly little is known about how the analytical parameters selected during these procedures (*e.g.,* read translation, homology classification thresholds, reference database) impact the accuracy of the resulting estimates of gene family abundance. This is especially problematic given that many of these methods lock users into specific parameters that may not be statistically appropriate for their data, which makes it hard to identify best practices in metagenome annotation.

We believe that identification of standardized approaches to metagenome annotation is best facilitated by analytically flexible, extensible, and stand-alone annotation software. While cloud-based and web-portal tools conveniently manage the entire annotation procedure (*i.e.,* from reads to gene family abundance estimates or functional profiles), and are widely used as a result, they have several limitations that result from their centralized design: (i) users tend to be limited to specific reference databases, (ii) it is difficult to extend the annotation software, and (iii) the requirement that users upload data to the server may present challenges of scale as the amount of metagenomic data increases. Hence, there is a growing need for stand-alone, open-source software that completely automates the annotation procedure. Most currently available stand-alone tools instead serve as add-ons to the core annotation procedure (*e.g*., HUMAnN, ShotgunFunctionalizeR [15]) or limit analyses to specific annotation parameters, such as the reference databases used to contextualize annotations (*e.g.,* MEGAN, RAMMCAP, SmashCommunity). In our assessment, there is a critical need for flexible and extensible software that automates shotgun metagenome annotation in a stand-alone environment and does so in a manner that is considerate of the statistical properties of the data.

We developed a new software tool, the Shotgun Metagenome Annotation Pipeline (ShotMAP), which is an end-to-end annotation workflow. To guide the development of ShotMAP, we conducted a large-scale simulation study and statistically assessed how the analytical parameters selected during metagenome annotation impact the accuracy of gene family abundance estimates. These simulations enabled us to identify a set of “best practices” that enable users to get the most out of their data given the read length of their metagenomes and the desired throughput. ShotMAP is analytically flexible to enable users to select settings appropriate for their data (*e.g.,* it is agnostic to the reference database used, mapping parameters are tunable to read length), and it can be implemented either on a local computer in a standalone environment or by interfacing with a Sun Grid Engine (SGE) or Portable Batch System (PBS) configured computing cluster.

We used ShotMAP in conjunction with analytical parameters that maximized annotation accuracy in our simulations to characterize the temporal genomic and transcriptomic variation of marine communities as well as the physiological variation of human gut microbiomes associated with inflammatory bowel disease. These analyses uncovered novel patterns of variation in photosynthetic protein families among marine communities and metabolic pathways that stratify Crohn’s disease-associated microbiomes from other patient populations.

## Results and Discussion

### Statistical simulations identify best practices in metagenome annotation

Quantifying and comparing the abundance of protein families across communities is critical for understanding how microbes have adapted to various environments [16] [17], and how variation in the functional composition of microbial communities can impact human health and disease [18]. However, it is not well understood how various bioinformatics procedures affect the accuracy of protein family abundance profiles.

Generally, metagenome annotation applies the following steps: (i) identification of putative protein coding sequences (*i.e.,* ORF prediction), (ii) comparison of predicted peptides to a protein family reference database using alignment algorithms, (iii) classification of predicted peptides into protein families (*i.e.,* homology designation), (iv) quantification of protein family abundance, and (v) comparison of differences in protein family profiles across samples. Different methods adopt varying parameters or specific procedures for each of these steps. For example, methods can vary in terms of how protein coding genes are predicted, the algorithms used to compare predicted peptides to the reference database, the thresholds used to identify homologs, and how protein family abundance should be quantified or normalized.

Here, we build upon previous work [7,19–22] and use mock communities and simulated metagenomes to systematically evaluate and optimize metagenome annotation [Fig. 1]. To our knowledge, our approach represents the first end-to-end evaluation and optimization of metagenome annotation. We specifically focus on identifying a set of best practices to maximize annotation accuracy while minimizing the computational time needed to generate these annotations. Furthermore, we pay special attention to the importance of read length and identify a number of read-length specific annotation parameters. Finally, we investigate the accuracy of both alpha and beta diversity estimation. An overview of how these simulations were performed and evaluated can be found in [Fig. 1], and the compositions of the simulated communities can be found in [S2 Figure] and [S3 Table].

**Figure 1.**
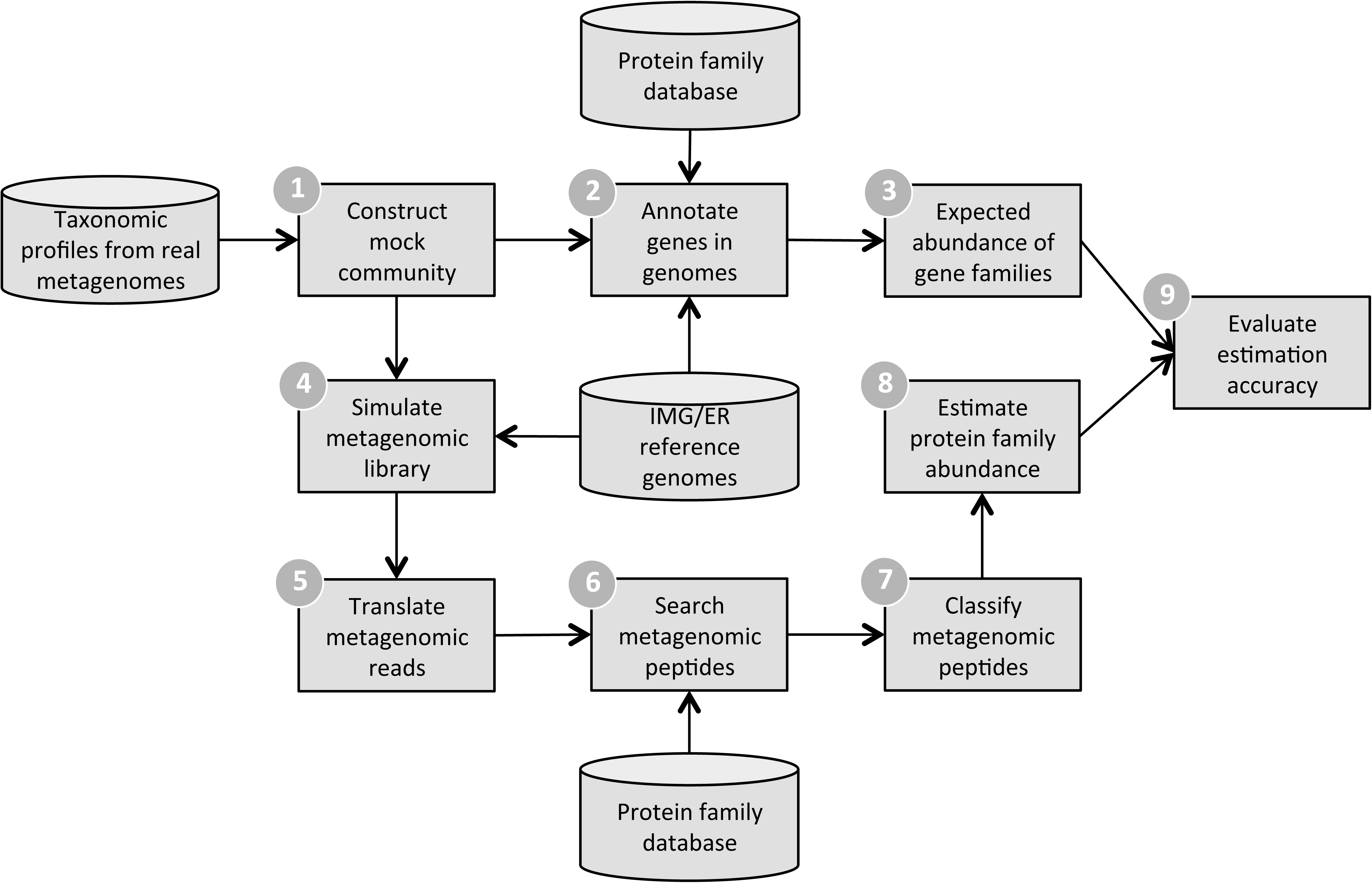
Metagenomic simulation framework. 1) Taxonomic profiles from real metagenomes are used to construct mock microbial communities. 2) Protein family annotations for SFams [23] are transferred to genes present in community members’ genomes. 3) The expected relative abundance of protein families is computed for the mock community. 4) Metagenomic reads are simulated from reference genomes that correspond to community members. 5) Simulated metagenomic reads are translated into predicted peptides. 6) Peptides are searched against the database of protein families used in (2) using the alignment tool RAPsearch2 [24] 7) Metagenomic peptides are classified into protein families according to their top-hit. Hits that do not satisfy the classification threshold are removed. 8) Classified metagenomic reads are used to estimate the relative abundance of protein families in the mock community. 9) Estimation accuracy is computed using an L1 distance between the expected and estimated relative abundance profiles.

### ORF prediction reduces data volume without sacrificing downstream accuracy

Read translation is the first step of many metagenome annotation pipelines. Naïve translation of metagenomic reads into all six possible reading frames (6FT) is commonly conducted, especially through tools like BLASTX [25]. While 6FT captures all true open reading frames (ORFs), it also increases the volume of query sequences and can lead to classification of spurious ORFs. Alternatively, *ab initio* metagenomic gene prediction [26], [27], [28] decreases the volume of queries, but can result in false-positive and false-negative reading frame predictions [29]. It is unclear which of these procedures results in more accurate functional profiles, and under what conditions. Previous work has evaluated the sensitivity and specificity of several metagenomic gene prediction tools [29], but it is not clear how these performance metrics translate into accuracy of protein family abundance estimates. Additionally, it is not clear how these tools compare to the widely used 6FT.

To address these questions, we used our statistical simulation pipeline [Fig. 1] to evaluate the accuracy of downstream functional abundance profiles and the volume of data that must be processed with different choices of translation method. Specifically, we evaluated performance for metagenomes from the mock community “160319967-stool1” using reads ranging from 50 bp to 3 kb simulated with a 1% uniform error rate. ORFs were searched and classified into SFams using RAPsearch2 while optimizing all other parameters (Methods). We ran 6FT in addition to three different metagenomic gene predictors – FragGeneScan [27], MetaGeneMark [28], and Prodigal [26] – and classified reads into protein families according to their top-hit across predicted ORFs (per-read annotation). Additionally, we evaluated a novel heuristic to rapidly filter short spurious ORFs [Fig. S4].

We found that the metagenomic gene finders reduced sequence volume by ∼85% relative to ORFs that had been naively translated in 6-frames [Fig. 2A], consistent with prior observations [29]. Despite this large decrease in sequence volume, there was relatively little decrease in accuracy compared to 6FT – particularly for metagenomes with reads at least 100 bp long [Fig. 2B]. For example, the use of Prodigal resulted in only 10.6% increase in L1 error relative to 6FT for 100-bp reads. We found similar results when using FragGeneScan, but found that MetaGeneMark performed significantly worse. The relative performance of all methods degraded with decreasing read length – Prodigal was the only tool that performed adequately using 70 bp reads and none of the tools performed adequately for read lengths shorter than 70 bp. These results indicate that metagenomic gene finders can be effectively used to reduce data volume for short-read metagenomes, but are only appropriate when read lengths exceed 70 bp. Additionally, 6FT resulted in the most accurate functional abundance profiles for short-read metagenomes, but resulted in ∼6x the number of initial sequences. Hence, we conclude that certain gene finders can significantly reduce data volume at a relatively low cost in accuracy for read lengths that are typical with current sequencing technologies.

**Figure 2:**
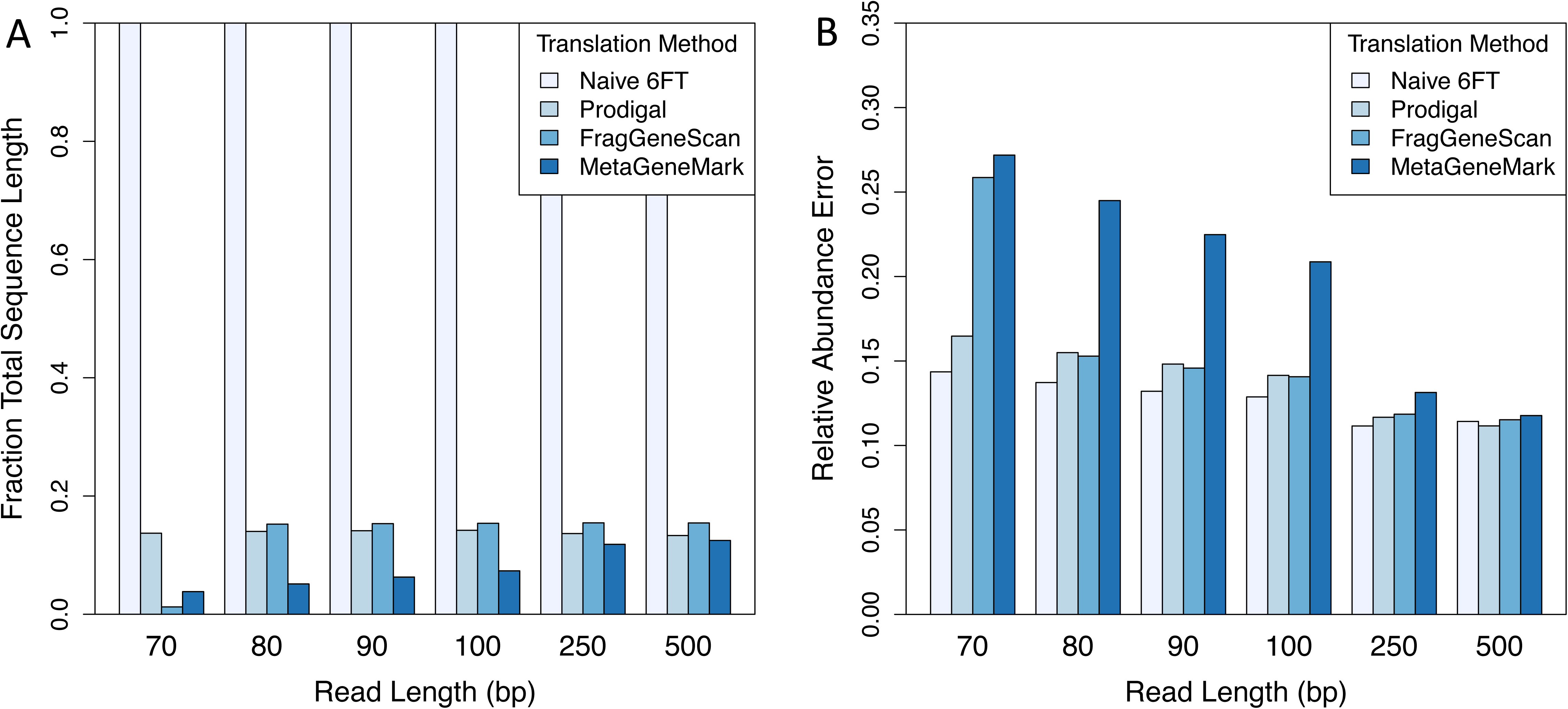
*Ab initio* gene prediction reduces data volume at a small cost to accuracy. Reads of various length (70 – 500 bp) were simulated with 1% error rate from mock community 160319967-stool1. Reads were translated naïvely in 6 frames (6FT) or were translated using a metagenomic gene prediction tool (Prodigal, FragGeneScan, or MetaGeneMark). Predicted ORFs were searched and classified into SFams using RAPsearch2. All other parameters were tuned to minimize relative abundance error. (A) Amino acid sequence length (relative to naïve 6FT) resulting from read translation. (B) Minimum relative abundance estimation error for mock community 160319967-stool1 corresponding to different translation methods.

### Annotating multiple ORFs per read is necessary with longer reads

In addition to the choice of translation method, it is important to consider how translated reads are annotated. One option is to classify a translated read according to the top-scoring hit across all of its predicted ORFs (i.e. one read, one annotation; referred to as “per-read” annotation). Alternatively, each predicted ORF can be classified independently (referred to as “per-ORF” annotation). Per-read annotation may make sense for short reads, which likely only contain a single true ORF, but may not be appropriate for longer reads that span multiple ORFs or contain overlapping ORFs.

Contrary to our expectations, we found that relative abundance error began to rapidly increase for reads longer than 500 bp, regardless of the translation method [Fig. 3A]. For example, metagenomes with 3,000 bp reads resulted in ∼3x more error than metagenomes with 250 bp reads. We hypothesized that this observation could be because longer reads contained multiple true ORFs that were not being annotated. To address this, we compared per-read and per-ORF annotation methods for long reads. Strikingly, we found that per-ORF annotation rescued performance for the 3-kb metagenomes and resulted in the most accurate functional abundance profiles across all read lengths [Fig. 3A]. Furthermore, when using per-ORF annotation, long reads actually benefitted from using Prodigal. We observed a clear switch in the optimal translation and annotation strategies at about 250 bp: metagenomes shorter than this benefitted from 6FT and per-read annotation, while metagenomes longer than this benefitted from Prodigal and per-ORF annotation [Fig. 3A]. We speculate these results are likely explained by (i) metagenomic gene-finders are less accurate for short-reads than for long-reads (26) and (ii) short reads usually only contain a single true ORF while long-reads are more likely to span multiple ORFs.

**Figure 3:**
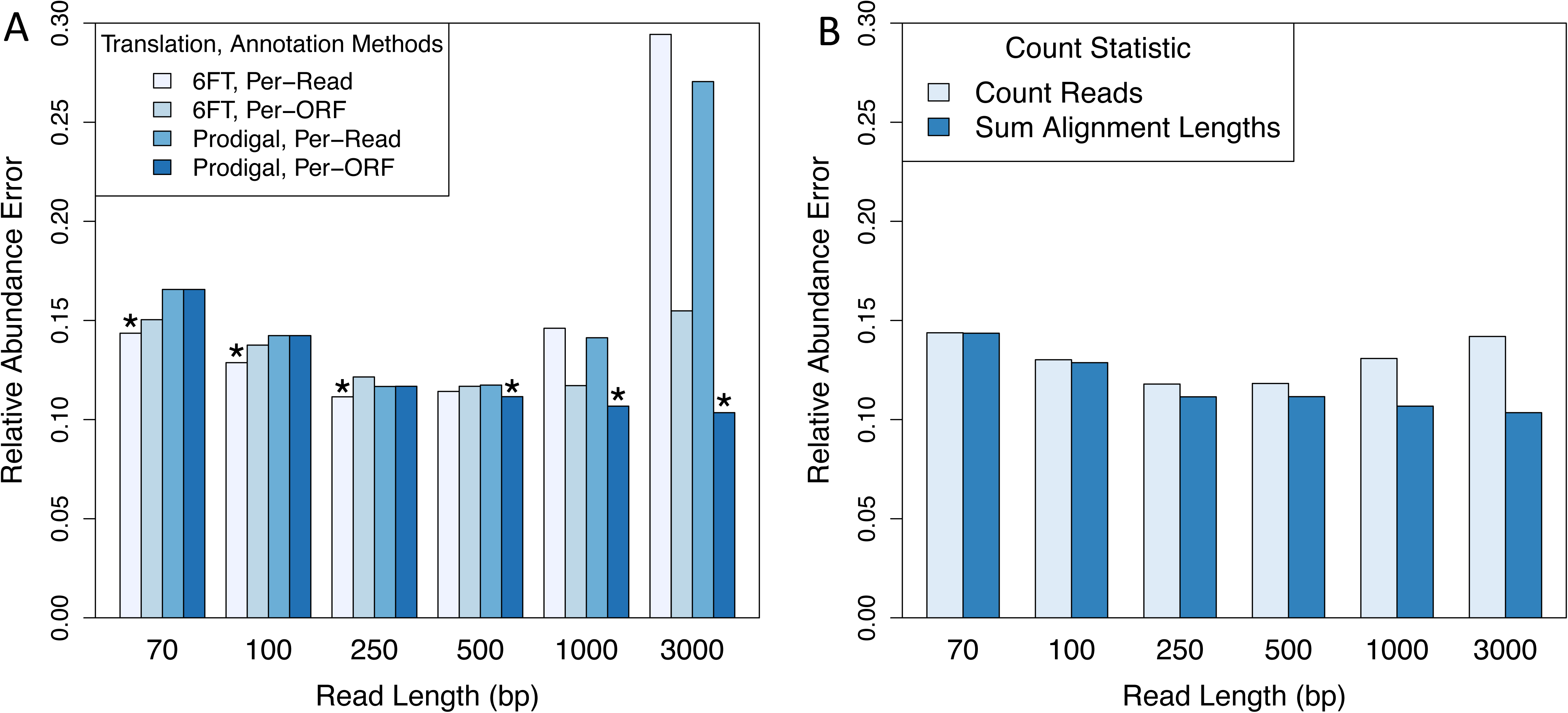
Short-reads and long-reads require different annotation strategies. Reads of various length (70 – 3,000 bp) were simulated with 1% error rate from mock community 160319967-stool1. (A) Predicted open reading frames (ORFs) were derived either via naïve 6- frame translation (6FT) or via the metagenomic gene finder Prodigal. Per-read annotation indicates that each read was classified according to the top-scoring hit across all of its ORFs. Per-ORF annotation indicates that each ORF was classified independently. Short reads benefit from 6FT and per-read annotation while long reads benefit from the gene finder Prodigal and per-ORF annotation. (B) Protein family abundances were estimated either by counting the number of hits to a family (count-based abundance) or by taking the sum of alignment lengths from hits (coverage-based abundance). In both cases, protein family abundance estimates were normalized by the gene length of reference sequences and scaled to sum to 1.0. The coverage-based abundance metric improves performance for long reads.

Finally, we were interested in exploring whether we could further increase performance for long-read metagenomes by accounting for reads that aligned at gene-boundaries. To address this, we evaluated two methods for computing family relative abundance. In the first, the abundance of each protein-coding gene family was obtained by counting all metagenomic reads classified into the family. We refer to this as count-based abundance. In the second method, instead of counting hits to a protein family, we counted the number of aligned residues across hits, which should account for reads that hang off the 5’ or 3’ end of a gene. We refer to this measure as coverage-based abundance. In both cases these metrics are normalized by gene length and converted to relative abundances that are scaled to sum to 1.0 (Methods). We found that both abundance metrics performed equivalently for short reads, but that coverage-based abundance significantly improved performance for reads longer than 500 bp [Fig. 3B]. For example, coverage-based abundance reduced relative abundance error from 0.14 to 0.1 (29% error reduction) for 3 kb reads.

In summary, naïve six-frame translation together with per-read annotation resulted in the most accurate estimates of protein family abundance for short-read metagenomes (≤250 bp), while *ab initio* gene prediction together with per-ORF annotation was optimal for reads longer than 250 bp. Additionally, we found that our coverage-based abundance metric was able to further increase accuracy for long-read metagenomes that presumably contain a greater proportion of reads that intersect gene boundaries. Overall, we found a good balance between algorithm speed, data reduction, and protein family abundance accuracy using the *ab initio* gene tool Prodigal. As a word of caution, while these methods have shown promise in our metagenomic simulations, it is not clear how they will perform in environments that contain organisms that utilize alternative genetic codes [30][31]. The performance of these methods in these kinds of environments needs to be investigated in greater detail in future work.

### Optimal alignment thresholds are read length specific and fail to reach commonly used E-value thresholds

After searching translated reads against a reference database, is it common practice to eliminate low-scoring or non-significant alignments and annotate each read according to its best hit in the reference database. While the choice of an E-value or bit-score threshold can have a major effect on the estimated relative abundance of protein families, there is little consensus and few guiding principles for choosing such a threshold. For example, over the past several years, published metagenomic studies have applied E-value thresholds ranging from 1e+1 [7] to 1e-10 [32] when annotating reads from Illumina and 454 sequencing technologies. Furthermore, because E-values depend on database size, read length, and the specific search tool used, it is not clear whether the same E-value threshold can be applied to the same effect across metagenomic studies. This is particularly relevant given the increasing size of reference databases and the increasing length of sequencing reads. Even when these variables are constant, different alignment tools and versions of BLAST can produce different E-values [33]. Here we used our simulation framework to systematically explore the effect these thresholds on the accuracy of metagenome annotation across simulated metagenomes with a wide range of different read lengths, sequencing error rates, and community compositions.

We began with an exploration of bit-score thresholds and found that (i) a precise bit-score threshold was critical to accurately estimate the relative abundance of protein families using short-read metagenomes [Fig. 4A], (ii) optimal bit-score thresholds were read length specific [Fig. 4A], and (iii) the bit-score thresholds we identified tended to correspond to non-significant E-values [S6 Figure]. Bit-score thresholds that were either too lenient or too stringent resulted in inaccurate estimates of protein family abundance, particularly for short-read metagenomes. For example, at 100 bp, we found that accuracy was maximized at a bit-score threshold of ∼31 bits; decreasing the threshold to 20 bits or increasing it to 50 bits increased error by 29-44%, which agrees with a previous report [34]. For reads longer than 100 bp, a precise bit-score threshold was not as important and similar accuracy was achieved over a wider range of thresholds [Fig. 4A], which is presumably due to an increased separation of false-positives from true-positives at longer read lengths. When applying optimal read-length thresholds it is important to recall that reads within a sample may vary in length, especially after trimming, so it may be desirable to use different thresholds for different reads (see below).

**Figure 4:**
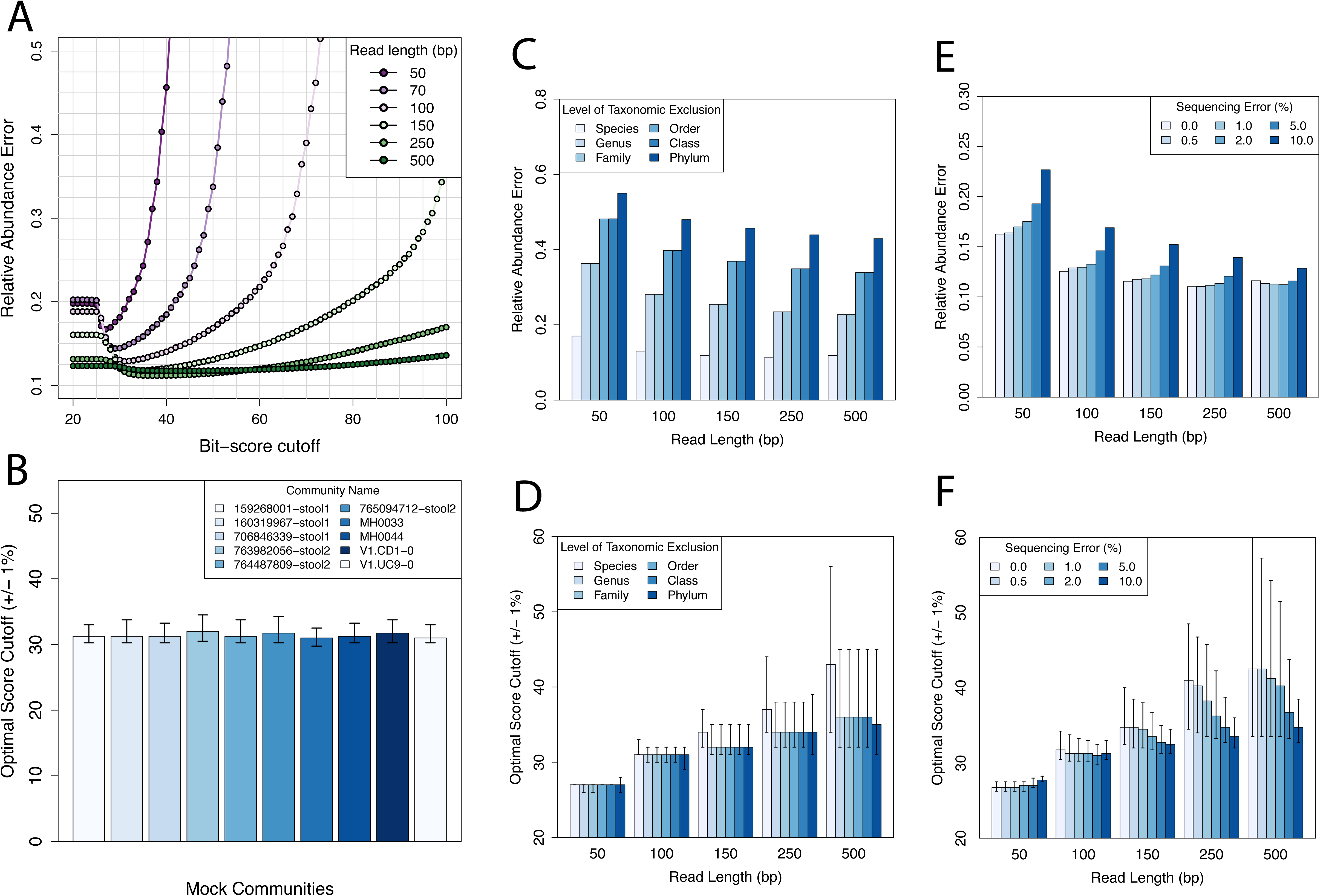
Relationship between read length, bit-score threshold, and prediction accuracy. (A) Simulated metagenomes (50 – 500 bp; 1% error rate; mock community 160319967-stool1) were searched and classified into SFams at different bit-score thresholds. At each threshold, L1 relative abundance error was calculated. (B) Simulated Illumina metagenomes from ten communities were searched and classified into SFams at different bit-score thresholds. Plotted is the optimal bit-score threshold for each community. Error bars indicate the range of bit-scores that result in L1 error within 1% of the optimal level. (C) Relative abundance error for simulated metagenomes of varying phylogenetic distance to reference genomes (50 – 500 bp; species to phylum taxonomic exclusion; 1% error rate; mock community 160319967-stool1). (D) Optimal bit-score thresholds error for metagenomes in (C). (E) Relative abundance error for simulated metagenomes of varying length and sequencing error (50 – 500 bp; 0-10% error rate; mock community 160319967-stool1). (F) Optimal bit-score thresholds for metagenomes in (E).

Next, we evaluated the stability of these optimal score cutoffs across different conditions. First, we repeated the experiment using 9 other mock communities and found that the optimal threshold for short-read metagenomes was quite stable regardless of community composition [Fig. 4B]. To further explore the effect of community composition, we performed clade exclusion experiments to simulate the presence of organisms from novel taxonomic groups (species, genera, families, orders, classes, phyla) in the metagenomes (Methods). We found that while we could accurately estimate protein-family abundance for communities composed of novel species, error quickly increased for communities composed of novel organisms at higher taxonomic levels (genera, families, order, classes, phyla) [Fig. 4C]. This result underscores the importance of using a phyolgenetically representative protein family database for a given study. While the phylogenetic novelty of the community had a major effect on the accuracy of functional profiles, it did not have a major effect on the optimal thresholds, which were generally stable [Fig. 4D]. In other words the optimal threshold is fairly constant across our clade-exclusion experiments, but the error associated with this threshold increases as the metagenome differs more from the database. Finally, we evaluated the effect of sequencing error rate. To address this, we simulated reads from mock community 160319967-stool1 using 50 to 500 bp reads that contained between 0 to 10% sequencing error. Interestingly, we found that error rates ≤2% had remarkable little effect on relative abundance error [Fig. 4E], indicating that homology searches are robust to typical error rates found in Illumina and 454 sequencing. Interestingly, the optimal score thresholds tended to decrease with an increasing error rate [Fig. 4F], which makes sense since homologs will end up with greater mismatches and lower scores.

In summary, we found that optimizing functional profiles from metagenomes required identifying read-length specific classification thresholds. These optimal thresholds were generally stable across different conditions (e.g. sequencing error, community composition, phylogenetic novelty) and generally resulted in low empirical false-positive rates for shuffled shotgun sequences. The importance of read-length specific alignment parameters has also been recognized for taxonomic annotation of metagenomes [35] and for estimation of average genome size from metagenomic data [36], consistent with our findings. Finally, while the cutoffs we identified maximized relative abundance accuracy in our simulations, these cutoffs may not be optimal for all protein families or for other types of downstream analyses.

### How much sequencing is enough: minimum number of reads for accurate estimates of physiological alpha-and beta-diversity

The choice of library size is a critical decision when designing a metagenomic experiment; insufficient sequencing depth can result in underestimates of diversity and high variance of abundance estimates, whereas extremely deep sequencing can be costly, time consuming, computationally challenging to analyze, and result in only marginal benefit. This decision is also relevant when reanalyzing data from published metagenomic studies, where analyzing a subset of reads may be sufficient for some purposes. Therefore, we investigated the role of sequencing depth on the ability to make accurate estimates of within (alpha) and between (beta) sample functional diversity. To our knowledge, this is the first systematic investigation into the role of sequencing depth on estimates of within and between sample functional diversity.

First, we determined the minimum number of reads for accurate estimates of within-sample protein family relative abundance. To address this question, we rarefied reads from our simulated 101 bp Illumina metagenomes, using between 10,000 and 1 million reads. At each sequencing depth, we used sampled reads to estimate the relative abundance of SFams, and compared these estimates to expected values using an L1 distance. As anticipated, we found that L1 error decreased with increasing sequencing depth and appeared to be close to an asymptote at a depth of ∼1 million reads [Fig. 5A]. While relative abundance error varied between mock communities, it appeared to reach an asymptote for all ten communities analyzed [Fig. 5A]. To investigate this further, we increased sequencing depth by two orders of magnitude to 100 million reads for one of the mock communities. With this massive increase in sequencing depth, there was only a marginal reduction in relative abundance error, from 0.13 to 0.11 [Fig. 5B].

**Figure 5:**
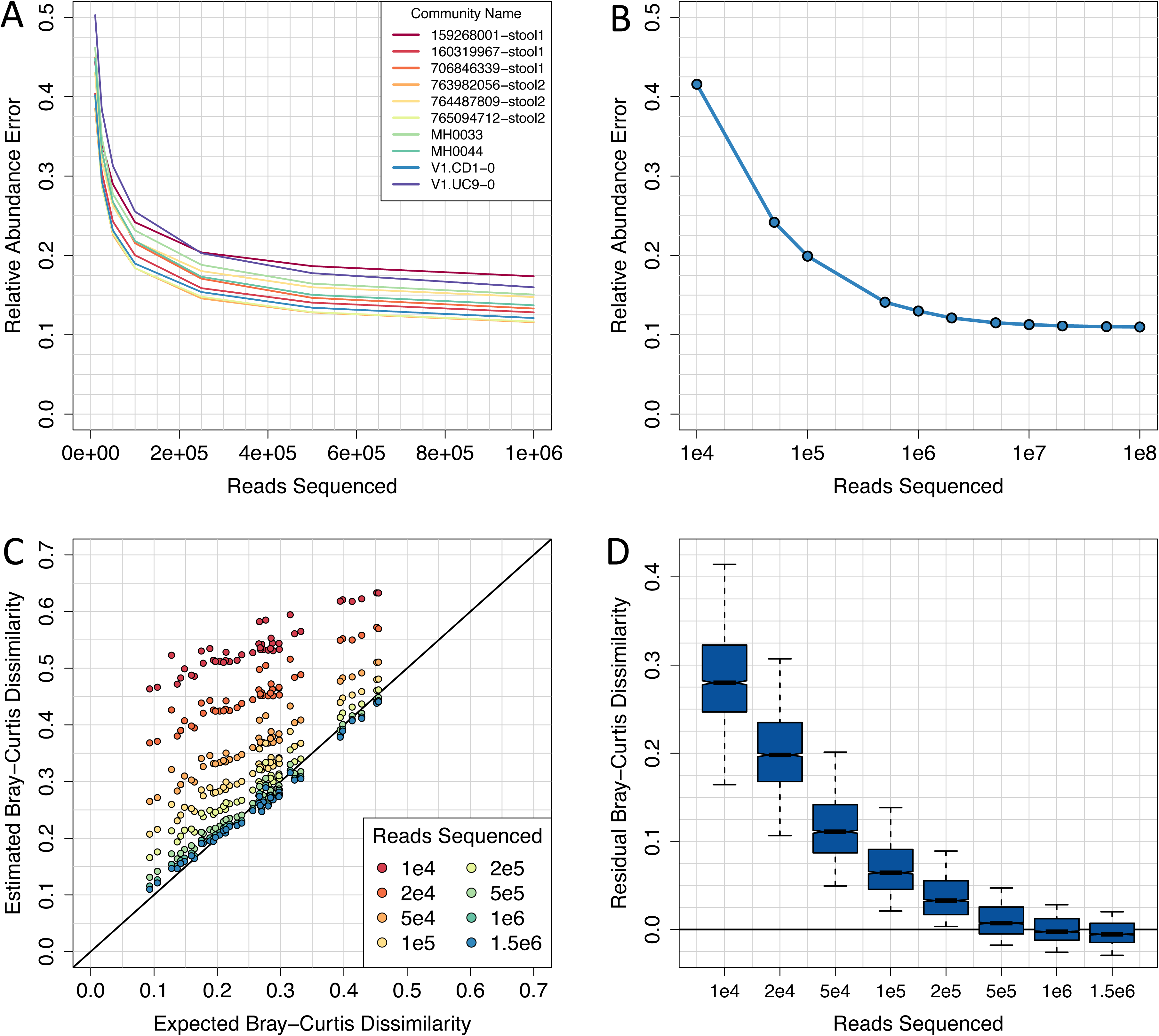
Shallow sequencing enables accurate estimates of alpha and beta functional diversity. (A) Relative abundance error for 101-bp Illumina metagenomes from 10 mock communities using between 10,000 and 1 million reads. (B) Relative abundance error for a 101-bp Illumina metagenomes from mock community 160319967-stool1 using between 10,000 and 100 million reads. (C) Expected versus observed functional distances for 10 mock communities using between 10,000 and 1.5 million 101-bp Illumina reads. (D) Distributions of Bray-Curtis dissimilarity error at each sequencing depth from (C).

Next, we assessed the effect of sequencing depth on the ability to accurately estimate functional distances between metagenomes (i.e. beta diversity). Like before, we used rarefied reads (10,000 to 1 million) to estimate the relative abundance of protein families from the ten mock communities and computed the Bray-Curtis dissimilarity between all pairs of samples (Methods). We found that Bray-Curtis dissimilarity could be accurately estimated using as few as 1 million reads (r2=0.99; mean residual distance = 0.01) [Fig. 5C-D]. At lower sequencing depths, functional distances were consistently overestimated, however the relative distances between pairs of samples was still highly correlated [Fig. 5C]. For example, at only 10,000 reads, Bray-Curtis dissimilarity was not accurately estimated (median residual distance = 0.28), but the relative distances between pairs of samples were still well correlated (r^2^ = 0.86).

While we found that 1 million reads was sufficient for overall estimates of alpha and beta functional diversity, it was not clear whether this was sufficient to accurately estimate the relative abundance of individual protein families. To investigate this, we bootstrapped reads from the 100 million read metagenome, using between 10,000 and 100 million reads. Based on this analysis, we found that protein families were Poisson distributed in the absence of biological variation, and that ∼100 classified reads/family was necessary for stable estimates of family abundance (i.e. coefficient of variation ≤ 0.10) [S7 Figure].

Together these results indicate that low-variance estimates of within and between sample functional diversity can be made using relatively small library sizes, but that deeper sequencing is required to accurately estimate the relative abundance of individual families. In the future, researchers may want to consider alternative studies designs in which many samples are sequenced at low sequencing depth. We speculate that such a design would be powered to detect major trends across large numbers of metagenomic samples.

## Accurate Metagenome Annotation Clarifies Community Functional Diversity and Identifies Biomarkers

### A novel Shotgun Metagenome Annotation Pipeline

Guided by the results of our simulation analyses, we developed extensible, open-source software to facilitate the accurate and automated inference of the biological functions encoded in a metagenome. This Shotgun Metagenome Annotation Pipeline (ShotMAP, https://github.com/sharpton/shotmap) is an analytical workflow that takes raw metagenomic reads as input and conducts ORF finding, homology inference, family classification, and uses robust statistical tests to evaluate physiological alpha-and beta-diversity [Fig. 4]. We include a complete description of the workflow in [S8 Text]. Our development of ShotMAP was born out of several observations. First, our statistical simulations indicate that the optimal analytical parameters for metagenome annotation depend on the context of the analysis (*e.g.,* read length), and most available tools lock users into particular settings (*e.g.,* E-value thresholds). ShotMAP, however, was designed to be analytically flexible, allowing users to select parameters appropriate for their investigation and data. For example, users can (1) select from a variety of gene-prediction and alignment algorithms, (2) tune the specific thresholds used to classify reads into families, and (3) apply a variety of family abundance normalizations (*e.g.,* average genome size, target family coverage, etc.). By default, ShotMAP will apply data-appropriate parameters (e.g., input data read-length dependent bit score threshold) that optimize the accuracy of estimating protein family abundance based on the results of our statistical simulations. Second, while different analyses may benefit from the use of a particular annotation space, most current tools only interface with specific protein family databases. We designed ShotMAP to be agnostic to the specific database used to conduct the annotation (including working with custom databases provided by the user) and to interface with either protein sequences or HMMER3-formatted HMMs [37]. Finally, ShotMAP fills the need for stand-alone, end-to-end annotation pipelines that start with unassembled reads and produce comparative inferences of functional diversity. While several cloud-based tools exist and are useful to the community (*e.g.,* MG-RAST), we anticipate that these resources will bottleneck analytical throughput as the number of metagenomes produced by the research community grows. Alternatively, ShotMAP can run on a multi-core benchtop computer and can optionally interface with an SGE-or PBS-configured cloud compute cluster. Notably, ShotMAP also allows users to manage their data and results through a MySQL relational database. It is open source to facilitate further development.

### Database decisions impact interpretation of marine metagenome seasonal variation

We leveraged ShotMAP’s ability to annotate metagenomes using different protein family databases to explore how database decisions impact estimates of functional diversity. We functionally annotated the Western English Channel L4 metagenomes and metatranscriptomes produced by Gilbert et al. [38] using three databases: KEGG Orthology groups (KOs), FIGfams [39], and SFams (Methods and [S9 Text]). We first compared the overall functional profiles of the L4 samples across databases by estimating protein family richness, Shannon entropy, and Good’s coverage. For both metagenomes and metatranscriptomes, we found that broad trends across samples were highly correlated between the three databases. For example, all databases produced protein family richness estimates that were largest in the January night and smallest in April day metagenomes [S10 Figure]. Additionally, we found that the functional diversity of metatranscriptomes was generally lower and more variable than that of the metagenomes, a finding consistent with those presented by Gilbert et al. that likely reflects a larger number of highly abundant families in the metatranscriptomic samples [S11 Figure]. However, we also found that specific measurements are impacted by the selection of a database. For example, KOs consistently produced lower diversity (*i.e.,* richness and Shannon entropy) and higher Good’s coverage estimates compared to the other databases. This is likely due to the order of magnitude smaller number of KOs. Additionally, SFams, the database containing the largest number of protein sequences, consistently classified more metagenomic reads than KOs and FIGfams [S12 Figure]. These analyses illustrate that the number of gene families and sequences in a protein family database systematically affect quantitative estimates of community diversity, but have less impact on global trends in diversity across samples.

Next, we sought to investigate the influence of the protein family database on inferences about specific gene families and functions. We first focused on photosynthesis, because Gilbert et al. observed that photosynthetic families were the most differentially abundant across seasons and between day and night [38]. We identified 141 KO’s and 84 FIGfams annotated as being involved in photosynthesis, and then compared normalized abundance estimates for each family across the metagenomic samples. Strikingly, reads were classified into 39 of the 141 KOs, but only eight of the 84 FIGfams [S13 Table]. Six of these eight FIGfams had homologs in the KO database, while the other two did not. Importantly, while these six families show a consistent pattern across samples, with relatively low abundance in summer and in the day as reported in Gilbert et al. [38], other photosynthetic KOs were equally abundant and showed opposite inter-sample patterns [S14 Figure]. Analysis of metatranscriptomes was qualitatively consistent with these observations, albeit limited by the low number of reads classified into photosynthetic families. Thus, despite having overall fewer gene families than FIGfams, the KO database has more annotated photosynthesis gene families and is therefore able to identify more reads as being involved in photosynthesis. This more diverse annotation reveals that the seasonal and diel trends observed with just six FIGfams were only part of a more complex pattern for genes with predicted roles in photosynthesis. Generalizing this result beyond photosynthesis, we observed seasonal and diel differences in a number of functional categories [S14 Figure], whereas Gilbert et al.’s analysis using FIGfams, only found differences in photosynthesis. We conclude that for individual functional categories, different protein family databases and annotation systems can influence data interpretation. Studies interested in robustly profiling specific functional categories would do well to consider annotating sequences with multiple protein family databases or annotation systems. The iterative classification option in ShotMAP makes this straightforward.

We used the summary statistics from ShotMAP to identify several additional aspects of the L4 datasets that heavily influence their interpretation (details in [S9 Text]). These analyses highlighted that both the number and proportion of reads mapped to low-abundance families are subject to stochastic effects (e.g., number of reads and classification rate per sample) and hence observed differences between samples may not be very robust.

### Metagenome annotation reveals functional variation associated with inflammatory bowel disease

We then used ShotMAP to determine if we could identify gene families whose abundance differs significantly between healthy and inflammatory bowel disease (IBD) microbiomes. We analyzed 43 stool metagenomes collected from a clinical population (the Metagenomic Species, or MGS, cohort) consisting of healthy controls (N=17), patients with Crohn’s Disease (CD; N=9), and patients with ulcerative colitis (UC; N=17) [40] to sensitively profile microbiome gene family diversity in each population. We used ShotMAP to classify 9M reads from each metagenome, which corresponded to the smallest number of reads generated per sample, into KOs (Methods). From this analysis, we found that CD associated microbiomes were physiologically distinct from healthy and UC associated microbiomes. Specifically, CD associated microbiomes exhibited lower protein family alpha-diversity as measured by richness, relative to microbiomes associated with either UC patients or healthy controls (p<0.05; [Fig. 7]), a finding that is consistent with prior observations [41–44]. Next, we used principal components analysis to cluster samples based on their protein family abundance profiles. Based on this analysis, we found evidence for differences in protein family abundance profiles between CD-associated microbiomes and their counterparts, based on the first and second principal components (together explaining 25.8% of the variation) and the first and third principal components (explaining 22.7% of the variation) [Fig. 7]. We also observe a moderate increase in the average genome size of the genomes that comprise CD microbiomes relative to the other population (p=0.1; [Fig. 7]). No evidence of substantial differentiation between UC-associated and healthy human microbiomes was identified. We additionally analyzed a separate IBD clinical cohort, the MetaHIT cohort (41), that similarly consisted of healthy controls (N=14), CD patients (N=4), and UC patients (N=21) (41). The differences that we observed between patient populations in this cohort were consistent with those in the MGS cohort, and the higher average genome size of CD microbiomes is significant in MetaHIT (p=0.005) [S16 Figure]. These data were additionally annotated using the MetaCyc and SFams databases, and the observed patterns were robust to differences in database selection.

**Figure 6:**
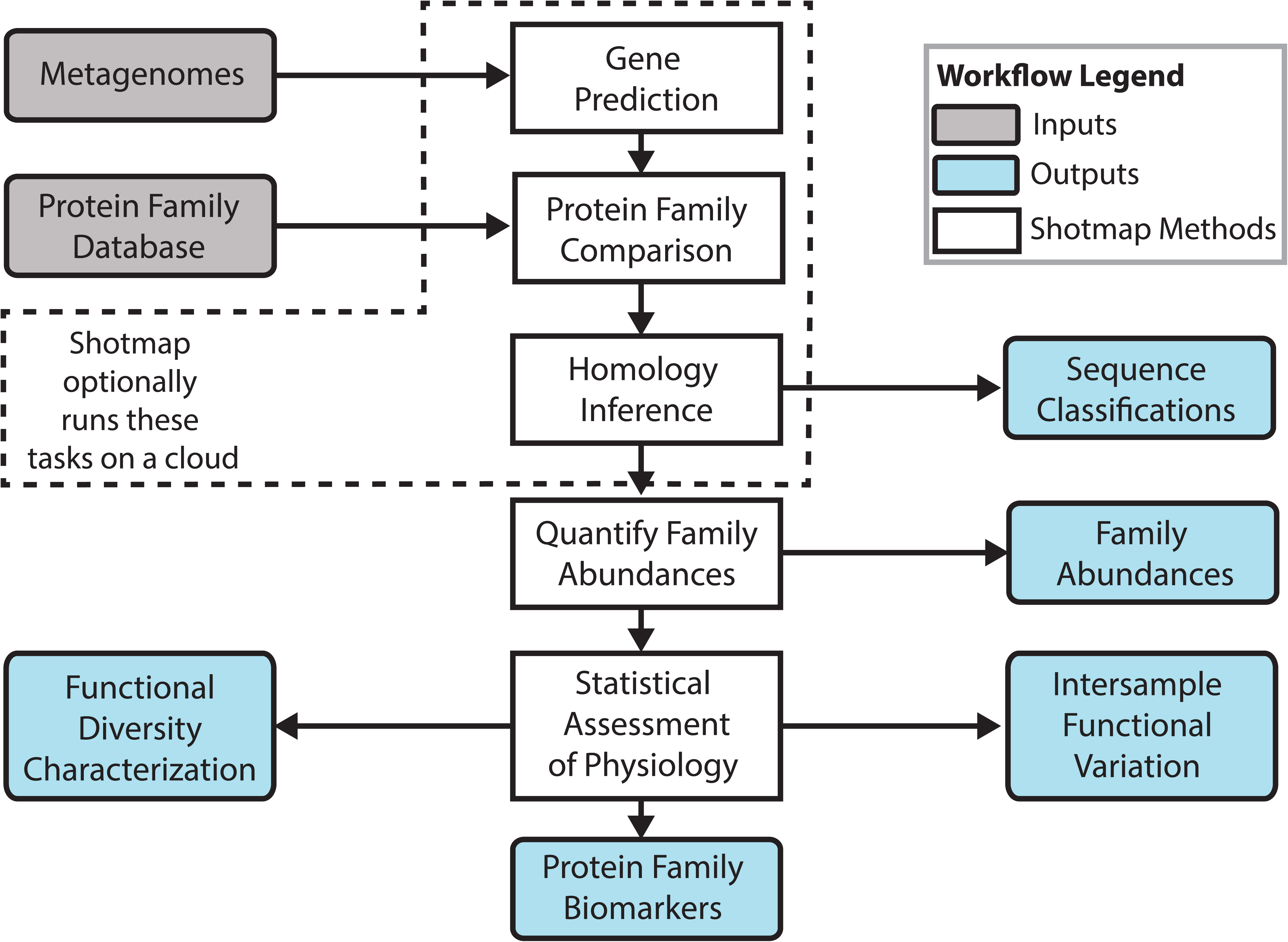
ShotMAP workflow. The Shotgun Metagenome Annotation Pipeline (ShotMAP) is an end-to-end metagenome annotation and analysis workflow. It takes as input metagenomic reads and a protein family database (grey boxes), and implements a variety of algorithms to predict genes, identify protein family homologs, quantify protein family abundances, and, optionally, statistically compare metagenomes (white boxes). ShotMAP produces a variety of outputs (blue boxes), including a mapping of sequences into protein families, an abundance profile of the protein families identified in the metagenome, estimates of protein family alpha-and beta-diversity, and a list of families that statistically stratify samples (*i.e.,* putative biomarkers). ShotMAP can be run on a local computer and can optionally interact with an SGE-configured cluster to manage computationally expensive tasks.

**Figure 7:**
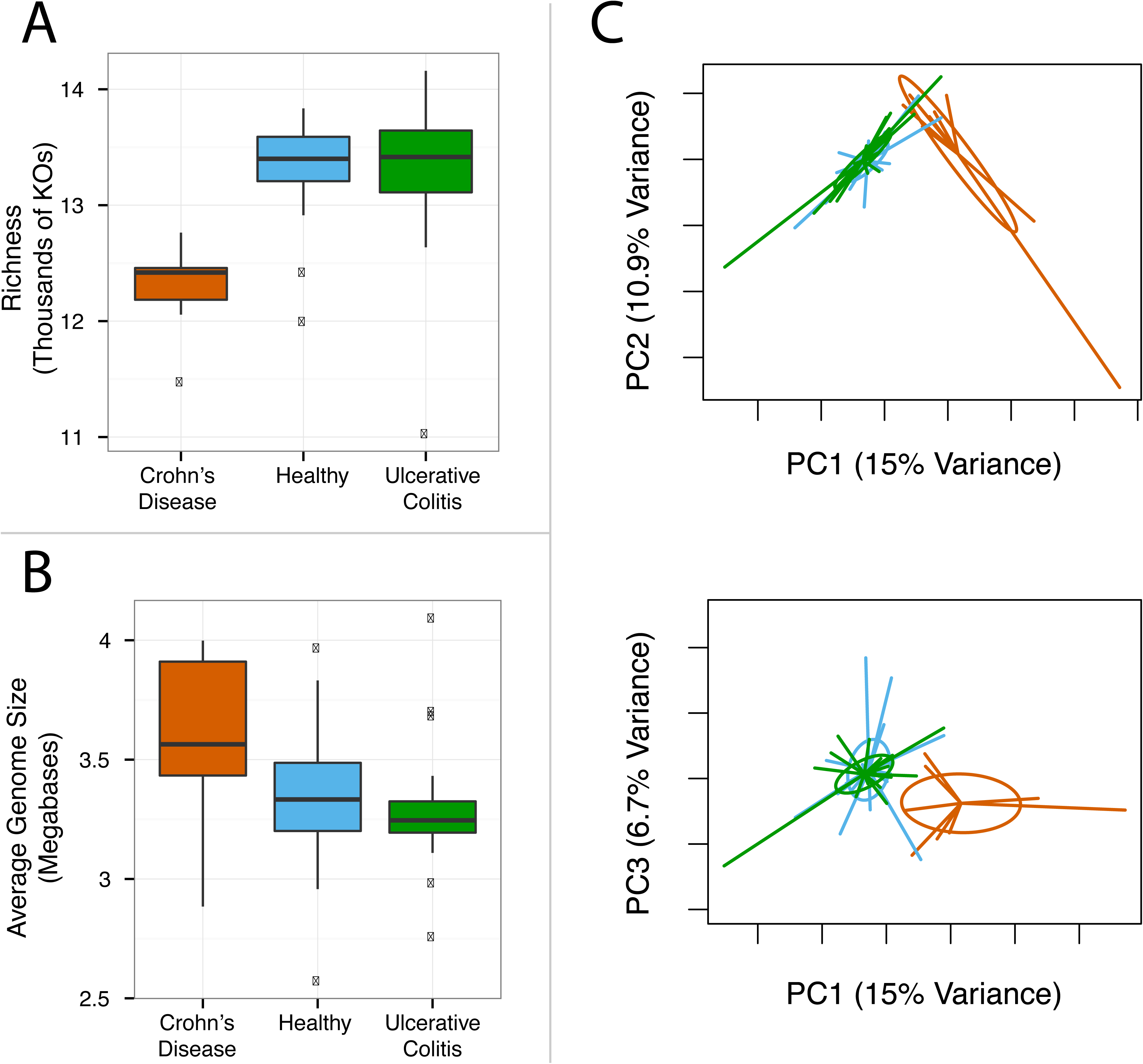
Crohn’s disease associated gut microbiomes are functionally distinct from microbiome associated with ulcerative colitis or healthy controls. Crohn’s disease (CD) associated gut microbiomes are functionally differentiated from their healthy and ulcerative colitis (UC) associated counterparts in the MGS cohort. (A) Boxplots illustrate that CD gut microbiomes exhibit lower protein family richness than UC and H microbiomes (p < 0.05). (B) The average genome size of the organisms in CD microbiomes is larger than the corresponding average in UC patients or healthy controls (p=0.1). (C) Principal components analysis (PCA) of protein family abundance profiles identifies differentiation in beta-diversity between CD (red) and non-CD populations (Ulcerative colitis, green; Healthy, blue) for the first two (upper panel) and first and third (lower panel) principal components. Here, PCA was conducted by scaling all KO abundances to have zero mean and unit variance, though similar structure is identified using different PCA parameters. Ellipses represent 95% confidence intervals. These trends are also observed in an independent analysis of the MetaHIT cohort [S16 Figure].

We then tested whether the abundances of any of the protein families detected by ShotMAP were associated with the clinical variables measured in the MetaHIT or MGS studies. These variables included gender, age, body mass index (BMI), and inflammatory bowel disease (IBD) status (none, ulcerative colitis (UC), and Crohn’s disease (CD)). In the MetaHIT dataset, we found few significant associations with age, gender, or body mass index, perhaps due in part to low power (Methods). In contrast, many KOs (796) were associated with IBD status (false discovery rate q <u>≤</u> 0.1). Interestingly, the gene families found to be significantly different tended to have altered abundance specifically in samples from patients with CD. When samples from UC and CD patients were grouped together, we found only two families that were differentially abundant between UC/CD and controls. These results were also seen in analysis of the larger MGS cohort: in fact, in the MGS dataset, no KEGG families had significant associations with gender, BMI, or even IBD when UC/CD were considered together, but we identified 3,511 differentially abundant KEGG families across disease groups when CD, UC, and control patients were considered separately.

These results are consistent with previous observations. Analysis of the taxonomic composition of MetaHIT gut microbiomes revealed that samples from UC patients were more similar to controls than to CD patients, which appeared dissimilar from both other groups [42]. Other research on separate cohorts also found that the community structure of fecal [45] and biopsy-associated microbiomes [46][47] differed substantially between UC and CD patients, with UC appearing more like controls; one study also saw similar trends in functional annotation from shallow shotgun sequencing [45]. UC and CD can also be differentiated serologically, with UC tending to be characterized more by autoantibodies directed at host neutrophils [48][49], though these may arise through cross-reactivity with microbial proteins like OmpC [50], and CD being characterized more by antibodies against cell-surface glycans [51], *Saccharomyces cerevisiae* [49], and proteins such as flagellin [52]. This difference in the antigens associated with CD vs. UC may relate in part to biomolecules made and presented by the gut flora.

To identify trends within these results, we used the MGS dataset, which provided the most power, to test whether CD-associated gene families were significantly enriched for any biological pathways of interest, using Fisher’s exact test. At a false discovery rate of q <u>≤</u> 0.25, we found 34 KEGG modules and 46 KEGG pathways that were elevated in patients with IBD [S17 Table]. One pair of KEGG modules that appeared to be more abundant in samples from patients with CD was “lipopolysaccharide [LPS] biosynthesis, KDO2-lipidA” (M00060;,q = 0.036) and “lipopolysaccharide biosynthesis, inner core => outer core => O-antigen” (M00080; q = 0.17). LPS (also called “endotoxin”), and in particular its component Lipid A, is known to modulate the innate immune system by binding to the toll-like receptor TLR4. Signaling through toll-like receptors such as TLR4 controls the release of pro-inflammatory cytokines (50). Notably, elevated antibodies against lipopolysaccharide have previously been linked to IBD and Crohn’s in particular (51)(52). Genome-wide association studies have also linked mutations in TLR4 to IBD in some (53), though not all cohorts (54)(55).

In the set of gene families more abundant in CD, we also observed that a KEGG module for the degradation of heparan sulfate (M00078; q = 0.121) appeared to be enriched, as well as the KEGG pathway “glycosaminoglycan degradation” (ko00531; q = 0.021) and a module for metabolism of a downstream metabolite, uronic acid (M00061; q = 0.12). Glycosaminoglycans (GAGs), of which heparan sulfate is one specific example, are associated with the mucosa of the GI tract and help to maintain its integrity. Prior work has observed that sulfated GAGs are depleted and disrupted in inflammatory bowel disease (56). Other studies have also found that gut flora, such as the commensal *B. thetaiotaomicron,* metabolize these GAGs, and that this metabolism contributes to their ability to colonize the gut (57)(58). The enrichment we observe may then suggest that flora associated with the Crohn’s-affected gut may be more likely to use these GAGs for nutrients than flora associated with either ulcerative colitis-affected guts or controls.

Finally, within the same set of genes, we also observe enrichments for KEGG modules and pathways describing the biosynthesis of several cofactors, including the B vitamins cobalamin (B_12_, M00122; q = 0.17), folate (B_9_, ko00790; q = 0.035), riboflavin (B_2_, ko00740; q = 0.16), pantothenate (B_5_, ko00770; q = 0.044), niacin (B_3_, ko00760; q = 0.035), biotin (B_7_, M00577; q = 0.17) and Vitamin K or menaquinone (M00116; q = 0.036). We also observed modules and pathways for the metabolism of the antioxidants Vitamin C (ascorbate, ko00053; q = 0.0014), in particular, its degradation (M00550; q = 0.16), and glutathione (ko00480; q = 0.052). Intriguingly, mucosal ascorbate levels in patients with IBD have been previously shown to be depleted [53] – a finding that was recapitulated in a mouse model [54]. Glutathione levels have also been shown to be depleted [55] particularly in inflamed CD mucosa [56]. Both ascorbate and glutathione are antioxidants; oxidative stress, including that initiated by immune cells, is recognized as playing a major role in causing mucosal injury in inflammatory bowel disease [57,58] and antioxidant levels have been linked to disease severity [59]. These results suggest the possibility that the microbiome, instead of being a totally independent factor in the progression of IBD, could exacerbate or otherwise modulate this important stressor.

One key function of B_12_ is to allow homocysteine to be converted into methionine. The increase in cobalamin biosynthetic genes we observe may translate into an increased ability to metabolize homocysteine, the levels of which have previously been shown to accumulate in patients with IBD (59-61). Indeed, a prior study [45] noticed that genes from gut microbiota of IBD patients were enriched for sulfur assimilation (which we also find here, 0.04 < q < 0.13, and which could also relate to metabolism of sulfated GAGs as above) and sulfur amino acid metabolism, which would include methionine. Alternatively, given that pyridoxine (61) and cobalamin (59) serum levels have been shown to be significantly lower in patients with IBD, and that in one study Crohn’s patients were observed to be more likely to be folate-or cobalamin-deficient than UC or control patients (62), bacteria that are capable of synthesizing their own B and K vitamins rather than scavenging from the environment could have an advantage within the IBD gut. Further investigation is necessary to determine how vitamin metabolism in Crohn’s patients affects and is affected by gut flora.

As a validation of these results, we also performed a similar analysis on the smaller MetaHIT cohort [42]. Looking at the set of gene families significantly increased in CD (q <u>≤</u> 0.1), we observed 16 enriched modules and 18 enriched pathways; of these, 7 (n. s.) and 16 (p = 1.3x10^−4^), respectively, were in common with the enrichments in the MGS dataset. Pathways enriched in common included vitamin metabolism (folate and biotin), GAG degradation, and glutathione metabolism; modules enriched in common included uronic acid metabolism, heparan sulfate degradation, and cobalamin biosynthesis. Finally, on the level of individual gene families, we found a substantial and significant overlap between gene families significantly associated with IBD status at an FDR of 0.1: after filtering out non-fully present KOs, 582 KOs were significantly associated in both out of 3,983 in either (p = 9.6x10^−6^). This agreement improved after considering only those KOs that were elevated in Crohn’s Disease (478 KOs in both out of 1,452 in either, p = 7.0x10^−9^). These results support our above findings and suggest that the Crohn’s-associated fecal microbiome may be distinguished from microbiota from healthy and Ulcerative Colitis patients in part by an increased abundance of gene families involved in vitamin, antioxidant, and glycosaminoglycan metabolism.

In summary, using ShotMAP, we characterized gene family abundance within the gut microbiota of the MGS cohort. This analysis showed that gut microbiota from CD patients had gene repertoires that were significantly different from UC patients and controls. Our results largely agree with analysis of a separate cohort (MetaHIT), which we present here, and also have features that overlap with prior investigations that considered a relatively limited number of individuals [44,60] or principally relied on ancestral state reconstruction using 16S rRNA sequences and validation through shallow metagenomic sequencing on a subset of samples [45,61]. For example, lipopolysaccharide biosynthetic genes were found to be elevated in IBD patients in several previous analyses [45,60,61], and one study of CD [44] found that ABC transporters were more abundant in CD metagenomes, which we also observe here (q = 1.2e-6). Furthermore, analysis of the the OSCCAR cohort (42), a separate group of Crohn’s cases and controls, found that not only LPS biosynthesis but GAG degradation, riboflavin biosynthesis, and sulfur metabolism were enriched in patients with ileal Crohn’s disease. In a complementary result, we found an increase in gene families involved in glutathione metabolism across two sets of CD patients, while Morgan et al. found that glutathione transport genes were more abundant in the microbiota of ileal CD patients.

Our results also had aspects that differed. For example, while we identified many metabolic pathways that increased in abundance in CD metagenomes, a separate study found that CD is principally associated with decreases in metabolic pathways and found very few CD-associated increases in pathway abundance [44]. Additionally, the analysis of the OSCCAR cohort identified a decrease in cobalamin biosynthetic genes among CD-associated metagenomes, while we observe increases in these genes among the corresponding populations in the cohorts studied here. One potential source of the observed differences between these cohorts may be explained by the fact we corrected for average genome size, which was found to be larger in CD metagenomes relative to controls. Additionally, Morgan et al. used whole genome sequences to estimate gene catalogs, and verified the results with low-coverage shotgun sequencing. In this verification step, the cobalamin biosynthetic genes were below the limit of detection. Another explanation could therefore be a difference between reference genomes and the particular strains represented in the OSCCAR cohort.

Further exploration of the metagenomic functional variation in larger and more diverse analyses will help clarify the ubiquity and robustness of the patterns observed here. These results validate the ability of ShotMAP to map gene family abundance in human microbiota, and demonstrate how gene family abundances, as well as groupings of genes into pathways, can be statistically associated with clinical variables.

## Conclusions

Our results indicate that the analytical decisions made during functional annotation of metagenomes can have a profound impact on the accuracy of estimating community function. Indeed, sequencing depth, read length, gene prediction and homology inference algorithms, and annotation database decisions can all impact the characterization of community function. Fortunately, we were also able to identify best-practices for most of the options that we explored (*e.g.,* classification score thresholds based on read lengths) and, in many cases, heuristics that balanced throughput with accuracy (*e.g.,* gene prediction with prodigal). While our simulations provide insight into the general statistical properties of metagenome annotation, the communities simulated here are based off of data obtained from human gut microbiome samples. The annotation prediction accuracy of metagenomes generated from very different types of communities may benefit from a similar simulation analysis.

By developing ShotMAP, an analytically flexible annotation workflow, we were able to reannotate previously published metagenomes and metatranscriptomes using the best practices identified in our simulation and identify novel physiological patterns and biomarkers. These analyses indicate that annotation decisions can impact the results and subsequent interpretation of the data. In a specific example, we were able to identify pathways enriched in metagenomes associated with Crohn’s disease-affected patients that provide insight into potential disease mechanisms and consequences (e.g., increased degradation of glycans). These findings may serve as clinically relevant diagnostic biomarkers, though additional study is needed to confirm these hypotheses.

## Materials and Methods

### Statistical assessment of metagenome annotation

#### Creation of *in silico* mock communities

We constructed mock microbial communities based on species abundance distributions observed in ten stool metagenomes from HMP [62] and MetaHIT [42] samples. Specifically, we leveraged work done by Schloissnig et al. [63], in which high quality reads from stool metagenomes were mapped to 909 non-redundant reference genomes. We determined the abundance of each genome to be the number of mapped reads to that genome divided by its genome size (*i.e.,* genome coverage). For all analyses, we used reference genome sequences downloaded from the IMG database of integrated microbial genomes [64] on March 28, 2013. Information on these communities can be found in [S2 Figure] and [S3 Table].

Protein coding sequences from each reference genome were annotated with gene families from the SFams v1 gene family database [23]. These annotations were used to determine the relative abundance of SFam gene families in each community. Specifically, the abundance of each gene family, *A*_*i*_, was defined as the copy number of the family in each genome, *C*_*i,j*_ for gene family *i* in genome *j),* weighted by the abundance of each genome in the community, 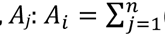 *(c*_*i,j*_ * *A*_*j*_). The relative abundance of each gene family, *R*_*i*_, was defined as the abundance normalized such that the sum of across families was equal to 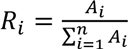.

#### Metagenome shotgun simulations

We used the shotgun sequence simulator Grinder [65] to simulate metagenomic libraries from the mock communities using IMG reference genomes [64]. In all libraries, the number of reads simulated from a genome was directly proportional to the relative abundance of that genome in the mock community. We used Grinder to generate a number of simulated shotgun libraries, which are indicated in the text and can be found online at: http://lighthouse.ucsf.edu/shotmapsims/

*Read-length simulations*: Here, we generated libraries of varying read length, from 25 to 3,000 base pairs (bp), from the mock community 160319967-stool1 [S3 Table]. Each library contained 1.3 million single-end reads, and were generated using a 1% uniform error rate and a 4:1 ratio of substitutions to indels, which is approximately consistent with raw error rates from Illumina sequencing platforms [66]. These libraries were used for the majority of experiments in this study.

*Ten-communities simulations:* Here we generated simulated Illumina libraries (Grinder options: -rd 101 -md poly4 3e-3 3.3e-8) from 10 mock communities [S2 Figure]. These libraries each contained ∼2 million single-end reads.

*Deep-sequenced simulation:* Here we generated a simulated Illumina library (Grinder options: -rd 101 -md poly4 3e-3 3.3e-8) from the mock community 160319967-stool1. This library contained ∼100 million single-end reads.

*Error-rate simulations:* Here, we generated libraries of varying error rate, from 0 to 10% from the mock community 160319967-stool1. Each library contained 1.3 million single-end reads, which varied in length from 50 to 500 bp.

*Error-model simulations:* Here, we simulated 100 bp single-end libraries of varying error models from the mock community 160319967-stool1. The error models included: Error-free (reads simulated without sequencing error); Uniform (1% uniform error rate); Illumina: (exponential error model); 454 (homopolymer error model); Sanger (linear error model ranging from 1% at the read start to 2% at the read end).

#### Read translation

We translated simulated DNA sequence reads into predicted open reading frames (ORFs) using three different approaches. In the first, we performed naïve translation of reads into all 6 possible reading frames (6FT) using the Transeq application from EMBOSS (v6.3.1) [67]. In the second approach, we took ORFs predicted from 6FT, split the sequences on stop-codons, and subsequently filtered these sequences to remove short spurious ORFs and reduce data volume [S4 Figure]. In the third approach, we used three popular tools – FragGeneScan [27], MetaGeneMark [28], and Prodigal [26] – to perform *ab initio* gene prediction. FragGeneScan (v1.18) was run using the options: -complete=0 -train=illumina_10; MetaGeneMark (v3.25) was run using the options: -m MetaGeneMark_v1.mod; Prodigal (v2.60) was run using the options: -p meta.

#### Homology search

We used RAPsearch2 (v2.15; default parameters) [24] to search predicted ORFs versus the SFams v1 protein family database v1 [23].

#### Clade exclusion

To simulate the presence of novel taxa in the mock communities, we held-back reference sequences belonging to organisms from the same taxonomic group as organisms in the metagenome, and classified reads into protein families using the remaining reference sequences. We performed this procedure and evaluated performance at the different taxonomic levels: strain, species, genus, family, order, class, and phylum. For example, at the genus level, this procedure would discard alignments between *Escherichia coli* shotgun sequences and all reference sequences from *Escherichia*; at the phylum level, this procedure would discard alignments between *E. coli* shotgun sequences and all reference sequences from Proteobacteria. Unless otherwise noted, clade exclusion was performed at the strain-level.

#### Classifying ORFs into protein families

We used two approaches for classifying predicted ORFs into protein families. In the first approach, we classified each metagenomic read according to the top-scoring hit across all of its predicted ORFs. By using this approach, each read can be annotated to only a single protein family. We refer to this as per-read annotation. In the second approach, we classified each predicted ORF independently. By using this approach, each read can be annotated to multiple protein families. We refer to as per-ORF annotation.

#### Relative abundance estimation

We used two metrics to estimate the abundance of protein families. In the first, the abundance of each protein-coding gene family, 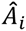, was obtained by counting all hits, *k,* to the family, *C*_*i*_ normalized by the target gene length of each hit, *L*_*k*_: 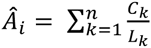. This normalization eliminates bias associated with gene length, whereby long genes would otherwise appear more abundant than short genes. We refer to this as count-based abundance. In the second method, instead of counting hits to a protein family, we counted the number of aligned residues across hits, *X*_*k*_which should account for reads that hang off the 5’ or 3’ end of a gene: 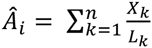. We refer to this measure as coverage-based abundance. In both cases the estimated relative abundance of each gene family, 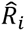, was normalized such that the sum of 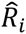 across families was equal to 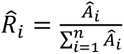.

#### Evaluating relative abundance error

We evaluated the prediction error for each community using the L1-distance between the expected, *R*_*i*_, and observed, 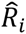, relative abundances of protein families. This metric is bounded by [0,1], a value of zero indicates no prediction error, and a value of 1.0 indicates the maximum prediction error: 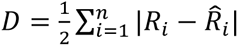.

### Analysis of Microbial Communities with ShotMAP

We used ShotMAP to analyze three previously published data sets, using parameters appropriate for the specific properties of the data, as identified by our statistical simulations [Results and SI]. First, we reprocessed the English Channel metagenomes (N=8) and metatranscriptomes (N=7) generated by Gilbert and colleagues [38] [S9 Text]. This study generated Roche/454 data from pelagic water samples taken from the L4 coastal ocean observatory site at both day and night time points in January, April and August of 2008. The metagenomes were annotated using six-frame translation and splitting on stop codons to predict ORFs, filtering ORFs less than 15 amino-acids in length, and using RAPsearch v2 to compare each ORF to protein sequences from three databases – SFams, KEGG, and FIGFams. Reads were determined to be homologs of the top-scoring family if they aligned to at least one of the family’s sequences with an alignment bit-score of at least 35. Metagenome and metatranscriptome family abundances were gene-length normalized to correct for variation in the length of families. Metagenome protein family abundances were normalized by their average genome size, as calculated by MicrobeCensus [36], to correct for variation in coverage across samples.

We then analyzed two sets of human stool metagenomes: one study produced by Qin and colleagues (41) as part of the MetaHIT clinical study of gut microbiomes associated with healthy (N=14), ulcerative colitis (N=21) and Crohn’s disease-affected (N=4) patients, and a later study produced by the MGS project, containing a distinct set of healthy (N = 17), ulcerative colitis (N = 17), and Crohn’s disease-affected patients (N = 9). The data was processed the same way as the L4 data, except that reads were mapped against the KEGG Orthology database and classified using minimum bit-score thresholds that optimized the accuracy of gene family abundance estimation based on each sample’s mean read length. We used the rank-based, non-parametric tests implemented in ShotMAP (i.e., Kruskal-Wallis tests for categorical variables and Kendall’s tau for continuous variables) to determine whether any protein families were associated with clinical variables measured by the MetaHIT authors.

Particularly in the case of the MetaHIT study, we found a large number of protein families with non-zero abundance in at least one sample. This degree of multiple hypothesis testing can make it difficult to control the false positive rate while retaining enough power to detect true positives. We therefore applied a statistical approach to filter gene families and reduce the number of tests [S15 Text]. We tested our approach by applying several different filters to the ShotMAP results, using statistics based on family abundance (*i.e.,* mean, standard deviation, coefficient of variation) as well as the fraction of samples in which the gene family was observed. We found that using only fully-observed gene families (i.e., those detected in all samples) improved the number of discoveries for each of the protein family databases tested. However, for the MGS data, filtering based on fully-observed gene families decreased the number of discoveries somewhat (3,511 vs. 4,926 with no filtering).

For each clinical variable, the per-protein family p-values were converted to q-values to correct for multiple testing using the procedure of Storey [68], setting a false discovery rate of 10%. We also tested whether significant protein families were enriched for particular biological pathways using Fisher’s exact test, again correcting for multiple testing and setting an FDR of 25%

## Acknowledgements

We are grateful to Aaron Darling, Guillaume Jospin, Morgan Langille, and Dongying Wu for their helpful discussions.

## Supporting Information Captions

**S1 Table. Current metagenome functional annotation and analysis tools.**

**S2 Figure: Species abundance distributions for ten mock microbial communities.** Relative abundances distributions for the top 20 most abundant species (out of 909 total) in each of the mock communities. Each community contains at least 500 species.

**S3 Table: Mock communities.** A list of the genomes and relative abundances present in each mock microbial community.

**S4 Figure: Filtering short spurious ORFs reduces data volume without increasing protein family abundance error.** Reads from simulated metagenomes (50-500 bp, 1% error rate, 1.2 million reads) were translated into all six possible reading frames and each reading frame was split into multiple peptides at stop codon positions. Peptides were filtered by their sequence length, and the remaining peptides were used to estimate the abundance of protein families in each metagenome. Each plot shows the results for a different metagenome (50, 100, 150, 250 and 500 bp reads). In each plot, the x-axis indicates the minimum length of peptides that were retained and used for homology interference and protein family abundance estimation. For example, a minimum ORF length of 40 indicates that peptides with less than 40 amino acids were discarded. The left y-axis and black curve indicate the fraction of total sequence length (in amino acids) that was left after filtering. The right y-axis and blue curve indicates the error in the resulting protein family abundance estimates.

**S5 Figure: Optimal score threshold for short-reads does not depend on sequencing error type**

Effect of the sequencing error model on (A) read-length specific cutoffs and (B) relative abundance error. All simulated reads were 100 bp long from community 160319967-stool1. Error-free: reads simulated without sequencing error; Uniform: 1% uniform error rate; Illumina: exponential error model; 454: homopolymer error model; Sanger: linear error model ranging from 1% at the read start to 2% at the read end.

**S6 Figure: Classification rates for randomly shuffled shotgun reads**

False positive rates for synthetic negatives. 500,000 reads were randomly sampled from community 160319967-stool1 at various read lengths (50, 100, 150, 250, 500). Each read was randomly shuffled in order to simulate synthetic negatives, which should not be classified into any protein family. Synthetic negatives were naively translated in 6 frames and searched against the SFams using RAPsearch2. Each read was classified according to its best hit across reading frames. (A-B) Empirical false positive rates at different bit-score cutoffs for different read lengths. Circles indicate optimal read-length specific cutoffs. (C) Empirical false positive rates versus E-values. Circles indicate optimal read-length specific cutoffs.

**S7 Figure: Effect of sequencing depth on individual families.** We used bootstrapping to estimate the variance of protein families as a function of the number of classified reads. Specifically, we resampled reads from the 100 million-read simulated metagenome, classified these reads into protein families, and computed abundance across families. We performed this procedure 100 times at each sequencing depth. a) In the absence of biological variation, protein families are Poisson distributed with mean equal to variance. b) The coefficient of variation (CV) of protein families decreases rapidly with increasing sequencing depth, and reaches an asymptote close to 1,000 reads and is sufficiently low at ∼100 reads. For example, families with ∼100 classified reads have a standard deviation of about 10 reads (CV=0.10). This indicates it would take ∼300,000 single-end Illumina reads to obtain a low-variance abundance estimate (CV ≤ 0.10) for a universally distributed single-copy gene present in a community with an average genome size of 3 Mb.

**S8 Text: Description of ShotMAP.**

**S9 Text: L4 Data Analysis Supplemental Methods.**

**S10 Figure: Broad diversity trends are consistent across databases used to annotate L4 samples.** Overall functional profiles for three databases calculated by ShotMAP. A-C Richness, Shannon entropy and Good’s coverage for KO, SFams and FIGfams over metagenomic samples for eight timepoints and metatranscriptomic samples for seven timepoints. Pearson’s rho pairwise across databases were (MG/MT): Richness KO:Sfams 0.67/0.99*; Sfams:FIGfams 0.87*/0.98*; KO:FIGfams 0.37/0.99*. Shannon KO:Sfams 0.47/0.93*;SFams:FIGfams 0.82*/0.89*;KO:FIGfams 0.39/0.93*.Good’s KO:SFams 0.92*/0.72; SFams:FIGfams 0.99*/0.94*;KO:FIGfams 0.91*/0.88*. \**p*<0.01

**S11 Figure: Metatranscriptomic samples have more highly abundant families than metagenomic samples from the same time points.** Boxplots for eight metagenomic and seven metatranscriptomic samples, which represent the distribution of the number of reads classified into each KO. Most KOs have a low classification rate, and metatranscriptomes (MT) are generally skewed towards having more highly abundant families than metagenomes (MG).

**S12 Figure: Protein family classification rates vary as a function of database, season, and data type.** The rate at which sequences are classified into protein families (classification rate, y-axis) varies across databases (different colored lines; KEGG Orthology Groups (KO), Sifting Families (SFams), FigFams (FF)), data type (metagenomes (MG), metatranscriptomes (MT)), and season (as indicated along the x-axis).

**S13 Table: Photosynthetic gene families detected in the L4 data.**

**S14 Figure: Inter-sample variation of photosynthetic gene families in the L4 data.** These plots illustrate change in the rarefied abundance (y-axis) across metagenomes (x-axis) of different KOs annotated as being involved in photosynthesis families. The upper plot represents the six KOs that show the same trends as Gilbert et al., which the lower plot represents six other KOs that show an opposite trend.

**S15 Text: MetaHIT Data Analysis Supplemental Methods.**

**S16 Figure: Protein family diversity differences between IBD populations in a smaller cohort (MetaHIT):** A smaller cohort of IBD patients and healthy controls (the MetaHIT cohort) were analyzed to assess the robustness of the patterns observed in the MGS cohort, which is described in the main text. (A) Boxplots illustrate that CD gut microbiomes exhibit lower protein family alpha-diversity than UC and H microbiomes (p < 0.05). Here, richness was calculated for each sample based on KO abundances, though the results are qualitatively consistent when SFams or MetaCyc families are quantified. (B) The average genome size of genomes comprising CD-associated microbiomes is significantly elevated relative to the other patient populations (p<0.05). (C) Principal components analysis (PCA) of protein family abundance profiles identifies differentiation in beta-diversity between CD (red) and non-CD populations. Here, PCA was conducted by zero centering and unit scaling KO abundances, though similar structure is identified using different protein family databases and PCA parameters. Ellipses represent 95% confidence intervals.

**S17 Table: Pathways enriched in the CD population.**

## References

1. Lozupone CA, Knight R. Global patterns in bacterial diversity. Proc Natl Acad Sci U S A [Internet]. 2007 Jul 3 [cited 2014 Dec 17];104(27):11436–40. Available from: http://www.pubmedcentral.nih.gov/articlerender.fcgi?artid=2040916&tool=pmcentrez&rendertype=abstract

2. Stegen JC, Lin X, Konopka AE, Fredrickson JK. Stochastic and deterministic assembly processes in subsurface microbial communities. ISME J [Internet]. 2012 Sep [cited 2015 Feb 18];6(9):1653–64. Available from: http://www.pubmedcentral.nih.gov/articlerender.fcgi?artid=3498916&tool=pmcentrez&rendertype=abstract

3. Degnan PH, Ochman H. Illumina-based analysis of microbial community diversity. ISME J [Internet]. International Society for Microbial Ecology; 2012 Jan [cited 2014 Mar 19];6(1):183–94. Available from: http://dx.doi.org/10.1038/ismej.2011.74

4. Lauber CL, Ramirez KS, Aanderud Z, Lennon J, Fierer N. Temporal variability in soil microbial communities across land-use types. ISME J [Internet]. 2013 Aug [cited 2014 Nov 23];7(8):1641–50. Available from: http://www.pubmedcentral.nih.gov/articlerender.fcgi?artid=3721119&tool=pmcentrez&rendertype=abstract

5. Sharpton TJ. An introduction to the analysis of shotgun metagenomic data. Front Plant Sci [Internet]. 2014 Jan [cited 2014 Jul 9];5:209. Available from: http://www.pubmedcentral.nih.gov/articlerender.fcgi?artid=4059276&tool=pmcentrez&rendertype=abstract

6. Huson DH, Auch AF, Qi J, Schuster SC. MEGAN analysis of metagenomic data. Genome Res [Internet]. 2007 Mar [cited 2014 Jul 9];17(3):377–86. Available from: http://www.pubmedcentral.nih.gov/articlerender.fcgi?artid=1800929&tool=pmcentrez&rendertype=abstract

7. Abubucker S, Segata N, Goll J, Schubert AM, Izard J, Cantarel BL, et al. Metabolic reconstruction for metagenomic data and its application to the human microbiome. PLoS Comput Biol [Internet]. 2012 Jan [cited 2014 Jan 23];8(6):e1002358. Available from: http://www.pubmedcentral.nih.gov/articlerender.fcgi?artid=3374609&tool=pmcentrez&rendertype=abstract

8. Li W. Analysis and comparison of very large metagenomes with fast clustering and functional annotation. BMC Bioinformatics [Internet]. 2009 Jan [cited 2014 Jan 25];10:359. Available from: http://www.pubmedcentral.nih.gov/articlerender.fcgi?artid=2774329&tool=pmcentrez&rendertype=abstract

9. Arumugam M, Harrington ED, Foerstner KU, Raes J, Bork P. SmashCommunity: a metagenomic annotation and analysis tool. Bioinformatics [Internet]. 2010 Dec 1 [cited 2014 Jan 23];26(23):2977–8. Available from: http://bioinformatics.oxfordjournals.org/content/26/23/2977.long

10. Kultima JR, Sunagawa S, Li J, Chen W, Chen H, Mende DR, et al. MOCAT: a metagenomics assembly and gene prediction toolkit. PLoS One [Internet]. Public Library of Science; 2012 Jan 17 [cited 2015 May 29];7(10):e47656. Available from: http://journals.PLoS.org/PLoSone/article?id=10.1371/journal.pone.0047656

11. Angiuoli S V, Matalka M, Gussman A, Galens K, Vangala M, Riley DR, et al. CloVR: a virtual machine for automated and portable sequence analysis from the desktop using cloud computing. BMC Bioinformatics [Internet]. 2011 Jan [cited 2012 Feb 29];12:356. Available from: http://www.pubmedcentral.nih.gov/articlerender.fcgi?artid=3228541&tool=pmcent rez&rendertype=abstract

12. Meyer F, Paarmann D, D’Souza M, Olson R, Glass EM, Kubal M, et al. The metagenomics RAST server - a public resource for the automatic phylogenetic and functional analysis of metagenomes. BMC Bioinformatics [Internet]. 2008 Jan [cited 2014 Jan 21];9:386. Available from: http://www.pubmedcentral.nih.gov/articlerender.fcgi?artid=2563014&tool=pmcentrez&rendertype=abstract

13. Dehal PS, Joachimiak MP, Price MN, Bates JT, Baumohl JK, Chivian D, et al. MicrobesOnline: an integrated portal for comparative and functional genomics. Nucleic Acids Res [Internet]. 2010 Jan [cited 2015 Feb 25];38(Database issue):D396–400. Available from: http://www.pubmedcentral.nih.gov/articlerender.fcgi?artid=2808868&tool=pmcentrez&rendertype=abstract

14. Markowitz VM, Chen I-MA, Chu K, Szeto E, Palaniappan K, Grechkin Y, et al. IMG/M: the integrated metagenome data management and comparative analysis system. Nucleic Acids Res [Internet]. 2012 Jan [cited 2015 Feb 25];40(Database issue):D123–9. Available from: http://www.pubmedcentral.nih.gov/articlerender.fcgi?artid=3245048&tool=pmcentrez&rendertype=abstract

15. Kristiansson E, Hugenholtz P, Dalevi D. Shotgun Functionalize R: an R-package for functional comparison of metagenomes. Bioinformatics [Internet]. 2009 Oct 15 [cited 2014 Jan 22];25(20):2737–8. Available from: http://bioinformatics.oxfordjournals.org/content/25/20/2737.abstract

16. Fierer N, Ladau J, Clemente JC, Leff JW, Owens SM, Pollard KS, et al. Reconstructing the microbial diversity and function of pre-agricultural tallgrass prairie soils in the United States. Science [Internet]. 2013 Nov 1 [cited 2014 Feb 20];342(6158):621–4. Available from: http://www.sciencemag.org/content/342/6158/621.full

17. HMP. A framework for human microbiome research. Nature [Internet]. NIH Public Access; 2012 Jun 14 [cited 2014 Jan 20];486(7402):215–21. Available from: http://europepmc.org/articles/PMC3377744/?report=abstract

18. Qin J, Li R, Raes J, Arumugam M, Burgdorf KS, Manichanh C, et al. A human gut microbial gene catalogue established by metagenomic sequencing. Nature [Internet]. 2010 Mar 4 [cited 2014 Jul 9];464(7285):59–65. Available from: http://www.pubmedcentral.nih.gov/articlerender.fcgi?artid=3779803&tool=pmcentrez&rendertype=abstract

19. Mavromatis K, Ivanova N, Barry K, Shapiro H, Goltsman E, McHardy AC, et al. Use of simulated data sets to evaluate the fidelity of metagenomic processing methods. Nat Methods [Internet]. 2007 Jun [cited 2014 Jan 24];4(6):495–500. Available from: http://www.ncbi.nlm.nih.gov/pubmed/17468765

20. Wommack KE, Bhavsar J, Ravel J. Metagenomics: read length matters. Appl Environ Microbiol [Internet]. 2008 Mar [cited 2015 Feb 28];74(5):1453–63. Available from: http://www.pubmedcentral.nih.gov/articlerender.fcgi?artid=2258652&tool=pmcentrez&rendertype=abstract

21. Dalevi D, Eriksen N. Expected Gene Order Distances and Model Selection in Bacteria. Bioinformatics [Internet]. 2008 Apr 1; Available from: http://bioinformatics.oxfordjournals.org/cgi/content/abstract/24/11/1332

22. Carr R, Borenstein E. Comparative analysis of functional metagenomic annotation and the mappability of short reads. PLoS One [Internet]. 2014 Jan [cited 2015 Feb 28];9(8):e105776. Available from: http://www.pubmedcentral.nih.gov/articlerender.fcgi?artid=4141809&tool=pmcent rez&rendertype=abstract

23. Sharpton TJ, Jospin G, Wu D, Langille MGI, Pollard KS, Eisen JA. Sifting through genomes with iterative-sequence clustering produces a large, phylogenetically diverse protein-family resource. BMC Bioinformatics [Internet]. 2012 Jan [cited 2014 Feb 14];13(1):264. Available from: http://www.biomedcentral.com/1471-2105/13/264

24. Zhao Y, Tang H, Ye Y. RAPSearch2: a fast and memory-efficient protein similarity search tool for next-generation sequencing data. Bioinformatics [Internet]. 2012 Jan 1 [cited 2014 Jan 21];28(1):125–6. Available from: http://www.pubmedcentral.nih.gov/articlerender.fcgi?artid=3244761&tool=pmcentrez&rendertype=abstract

25. Altschul SF, Gish W, Miller W, Myers EW, Lipman DJ. Basic local alignment search tool. J Mol Biol [Internet]. 1990 Oct 5 [cited 2014 Jul 10];215(3):403–10. Available from: http://www.ncbi.nlm.nih.gov/pubmed/2231712

26. Hyatt D, LoCascio PF, Hauser LJ, Uberbacher EC. Gene and translation initiation site prediction in metagenomic sequences. Bioinformatics [Internet]. 2012 Sep 1 [cited 2015 Feb 25];28(17):2223–30. Available from: http://www.ncbi.nlm.nih.gov/pubmed/22796954

27. Rho M, Tang H, Ye Y. FragGeneScan: predicting genes in short and error-prone reads. Nucleic Acids Res [Internet]. 2010 Nov [cited 2014 Jan 30];38(20):e191. Available from: http://www.pubmedcentral.nih.gov/articlerender.fcgi?artid=2978382&tool=pmcentrez&rendertype=abstract

28. Zhu W, Lomsadze A, Borodovsky M. Ab initio gene identification in metagenomic sequences. Nucleic Acids Res [Internet]. 2010 Jul 1 [cited 2014 Jan 23];38(12):e132. Available from: http://nar.oxfordjournals.org/content/38/12/e132.abstract

29. Trimble WL, Keegan KP, D’Souza M, Wilke A, Wilkening J, Gilbert J, et al. Short-read reading-frame predictors are not created equal: sequence error causes loss of signal. BMC Bioinformatics [Internet]. 2012 Jan [cited 2014 Jan 28];13(1):183. Available from: http://www.biomedcentral.com/1471-2105/13/183

30. McCutcheon JP, McDonald BR, Moran NA. Origin of an alternative genetic code in the extremely small and GC-rich genome of a bacterial symbiont. PLoS Genet [Internet]. 2009 Jul [cited 2015 Feb 25];5(7):e1000565. Available from: http://www.pubmedcentral.nih.gov/articlerender.fcgi?artid=2704378&tool=pmcentrez&rendertype=abstract

31. Bertram G, Innes S, Minella O, Richardson J, Stansfield I. Endless possibilities: translation termination and stop codon recognition. Microbiology [Internet]. 2001 Feb [cited 2015 Feb 25];147(Pt 2):255–69. Available from: http://www.ncbi.nlm.nih.gov/pubmed/11158343

32. Fierer N, Ladau J, Clemente JC, Leff JW, Owens SM, Pollard KS, et al. Reconstructing the microbial diversity and function of pre-agricultural tallgrass prairie soils in the United States. Science [Internet]. 2013 Nov 1 [cited 2014 Feb 20];342(6158):621–4. Available from: http://www.sciencemag.org/content/342/6158/621.abstract

33. Hauswedell H, Singer J, Reinert K. Lambda: the local aligner for massive biological data. Bioinformatics [Internet]. 2014 Sep 1 [cited 2015 Feb 25];30(17):i349–55. Available from: http://www.pubmedcentral.nih.gov/articlerender.fcgi?artid=4147892&tool=pmcentrez&rendertype=abstract

34. Du R, Mercante D, Fang Z. An artificial functional family filter in homolog searching in next-generation sequencing metagenomics. PLoS One [Internet]. 2013 Jan [cited 2015 Feb 25];8(3):e58669. Available from: http://www.pubmedcentral.nih.gov/articlerender.fcgi?artid=3597637&tool=pmcentrez&rendertype=abstract

35. Liu B, Gibbons T, Ghodsi M, Treangen T, Pop M. Accurate and fast estimation of taxonomic profiles from metagenomic shotgun sequences. BMC Genomics [Internet]. 2011 Jan [cited 2014 Jan 22];12 Suppl 2(Suppl 2):S4. Available from: http://www.biomedcentral.com/1471-2164/12/S2/S4

36. Nayfach S, Pollard KS. Average genome size estimation enables accurate quantification of gene family abundance and sheds light on the functional ecology of the human microbiome [Internet]. bioRxiv. Cold Spring Harbor Labs Journals; 2014 Sep [cited 2015 Feb 25]. Available from: http://biorxiv.org/content/early/2014/09/11/009001.abstract

37. Eddy SR. Accelerated Profile HMM Searches. PLoS Comput Biol [Internet]. 2011 Oct [cited 2014 Jul 16];7(10):e1002195. Available from: http://www.pubmedcentral.nih.gov/articlerender.fcgi?artid=3197634&tool=pmcentrez&rendertype=abstract

38. Gilbert JA, Field D, Swift P, Thomas S, Cummings D, Temperton B, et al. The taxonomic and functional diversity of microbes at a temperate coastal site: a “multiomic” study of seasonal and diel temporal variation. Rodriguez-Valera F, editor. PLoS One [Internet]. Public Library of Science;. 2010 Jan [cited 2013 May 21];5(11):e15545. Available from: http://dx.PLoS.org/10.1371/journal.pone.0015545

39. Meyer F, Overbeek R, Rodriguez A. FIGfams: yet another set of protein families. Nucleic Acids Res [Internet]. 2009 Nov [cited 2015 Jan 6];37(20):6643–54. Available from: http://www.pubmedcentral.nih.gov/articlerender.fcgi?artid=2777423&tool=pmcentrez&rendertype=abstract

40. Nielsen HB, Almeida M, Juncker AS, Rasmussen S, Li J, Sunagawa S, et al. Identification and assembly of genomes and genetic elements in complex metagenomic samples without using reference genomes. Nat Biotechnol [Internet]. Nature Publishing Group; 2014 Jul 6 [cited 2014 Jul 9];32(8):822–8. Available from: http://www.nature.com/nbt/journal/v32/n8/full/nbt.2939.html#supplementary-information

41. Manichanh C, Rigottier-Gois L, Bonnaud E, Gloux K, Pelletier E, Frangeul L, et al. Reduced diversity of faecal microbiota in Crohn’s disease revealed by a metagenomic approach. Gut [Internet]. 2006 Feb 1 [cited 2012 Oct 4];55(2):205–11. Available from: http://gut.bmj.com/content/55/2/205

42. Qin J, Li R, Raes J, Arumugam M, Burgdorf KS, Manichanh C, et al. A human gut microbial gene catalogue established by metagenomic sequencing. Nature. Nature Publishing Group; 2010 Mar 4;464(7285):59–65.

43. Lepage P, Häsler R, Spehlmann ME, Rehman A, Zvirbliene A, Begun A, et al. Twin Study Indicates Loss of Interaction Between Microbiota and Mucosa of Patients With Ulcerative Colitis. Gastroenterology [Internet]. 2011 Jul [cited 2015 Apr 3];141(1):227–36. Available from: http://www.ncbi.nlm.nih.gov/pubmed/21621540

44. Erickson AR, Cantarel BL, Lamendella R, Darzi Y, Mongodin EF, Pan C, et al. Integrated metagenomics/metaproteomics reveals human host-microbiota signatures of Crohn’s disease. PLoS One [Internet]. Public Library of Science; 2012 Jan 28 [cited 2015 Jun 22];7(11):e49138. Available from: http://journals.PLoS.org/PLoSone/article?id=10.1371/journal.pone.0049138

45. Morgan XC, Tickle TL, Sokol H, Gevers D, Devaney KL, Ward D V, et al. Dysfunction of the intestinal microbiome in inflammatory bowel disease and treatment. Genome Biol [Internet]. 2012 Jan [cited 2014 Jan 23];13(9):R79. Available from: http://genomebiology.com/2012/13/9/R79

46. Gophna U, Sommerfeld K, Gophna S, Doolittle WF, Veldhuyzen van Zanten SJO. Differences between tissue-associated intestinal microfloras of patients with Crohn’s disease and ulcerative colitis. J Clin Microbiol [Internet]. 2006 Nov [cited 2015 Feb 26];44(11):4136–41. Available from: http://www.pubmedcentral.nih.gov/articlerender.fcgi?artid=1698347&tool=pmcentrez&rendertype=abstract

47. Hansen R, Russell RK, Reiff C, Louis P, McIntosh F, Berry SH, et al. Microbiota of de-novo pediatric IBD: increased Faecalibacterium prausnitzii and reduced bacterial diversity in Crohn’s but not in ulcerative colitis. Am J Gastroenterol [Internet]. 2012 Dec [cited 2015 Feb 26];107(12):1913–22. Available from: http://www.ncbi.nlm.nih.gov/pubmed/23044767

48. Saxon A, Shanahan F, Landers C, Ganz T, Targan S. A distinct subset of antineutrophil cytoplasmic antibodies is associated with inflammatory bowel disease. J Allergy Clin Immunol [Internet]. 1990 Aug [cited 2015 Feb 26];86(2):202–10. Available from: http://www.ncbi.nlm.nih.gov/pubmed/2200820

49. Joossens S, Reinisch W, Vermeire S, Sendid B, Poulain D, Peeters M, et al. The value of serologic markers in indeterminate colitis: a prospective follow-up study. Gastroenterology [Internet]. 2002 May [cited 2015 Feb 26];122(5):1242–7. Available from: http://www.ncbi.nlm.nih.gov/pubmed/11984510

50. Cohavy O, Bruckner D, Gordon LK, Misra R, Wei B, Eggena ME, et al. Colonic bacteria express an ulcerative colitis pANCA-related protein epitope. Infect Immun [Internet]. 2000 Mar [cited 2015 Feb 26];68(3):1542–8. Available from: http://www.pubmedcentral.nih.gov/articlerender.fcgi?artid=97313&tool=pmcentrez&rendertype=abstract

51. Dotan I, Fishman S, Dgani Y, Schwartz M, Karban A, Lerner A, et al. Antibodies against laminaribioside and chitobioside are novel serologic markers in Crohn’s disease. Gastroenterology [Internet]. 2006 Aug [cited 2015 Feb 26];131(2):366–78. Available from: http://www.ncbi.nlm.nih.gov/pubmed/16890590

52. Lodes MJ, Cong Y, Elson CO, Mohamath R, Landers CJ, Targan SR, et al. Bacterial flagellin is a dominant antigen in Crohn disease. J Clin Invest [Internet]. 2004 May [cited 2015 Feb 26];113(9):1296–306. Available from: http://www.pubmedcentral.nih.gov/articlerender.fcgi?artid=398429&tool=pmcentrez&rendertype=abstract

53. Buffinton GD, Doe WF. Altered ascorbic acid status in the mucosa from inflammatory bowel disease patients. Free Radic Res [Internet]. 1995 Feb [cited 2015 Jun 19];22(2):131–43. Available from: http://www.ncbi.nlm.nih.gov/pubmed/7704184

54. Blackburn AC, Doe WF, Buffinton GD. Colonic antioxidant status in dextran sulfate-induced colitis in mice. Inflamm Bowel Dis [Internet]. 1997 Jan [cited 2015 Jun 19];3(3):198–203. Available from: http://www.ncbi.nlm.nih.gov/pubmed/23282805

55. Miralles-Barrachina O, Savoye G, Belmonte-Zalar L, Hochain P, Ducrotté P, Hecketsweiler B, et al. Low levels of glutathione in endoscopic biopsies of patients with Crohn’s colitis: the role of malnutrition. Clin Nutr [Internet]. 1999 Oct [cited 2015 Jun 19];18(5):313–7. Available from: http://www.ncbi.nlm.nih.gov/pubmed/10601540

56. Pinto MAS, Lopes MS-MS, Bastos STO, Reigada CLL, Dantas RF, Neto JCB, et al. Does active Crohn’s disease have decreased intestinal antioxidant capacity? J Crohns Colitis [Internet]. 2013 Oct [cited 2015 Jun 19];7(9):e358–66. Available from: http://www.ncbi.nlm.nih.gov/pubmed/23523266

57. Grisham MB. Oxidants and free radicals in inflammatory bowel disease. Lancet [Internet]. 1994 Sep 24 [cited 2015 Jun 14];344(8926):859–61. Available from: http://www.ncbi.nlm.nih.gov/pubmed/7916405

58. Bhattacharyya A, Chattopadhyay R, Mitra S, Crowe SE. Oxidative stress: an essential factor in the pathogenesis of gastrointestinal mucosal diseases. Physiol Rev [Internet]. 2014 Apr [cited 2015 Jun 19];94(2):329–54. Available from: http://www.pubmedcentral.nih.gov/articlerender.fcgi?artid=4044300&tool=pmcentrez&rendertype=abstract

59. Hengstermann S, Valentini L, Schaper L, Buning C, Koernicke T, Maritschnegg M, et al. Altered status of antioxidant vitamins and fatty acids in patients with inactive inflammatory bowel disease. Clin Nutr [Internet]. 2008 Aug [cited 2015 Jun 24];27(4):571–8. Available from: http://www.ncbi.nlm.nih.gov/pubmed/18316141

60. Greenblum S, Turnbaugh PJ, Borenstein E. Metagenomic systems biology of the human gut microbiome reveals topological shifts associated with obesity and inflammatory bowel disease. Proc Natl Acad Sci U S A [Internet]. 2012 Jan 10 [cited 2014 Nov 7];109(2):594–9. Available from: http://www.pnas.org/content/109/2/594.full

61. Gevers D, Kugathasan S, Denson LA, Vázquez-Baeza Y, Van Treuren W, Ren B, et al. The treatment-naive microbiome in new-onset Crohn’s disease. Cell Host Microbe [Internet]. 2014 Mar 12 [cited 2014 Jul 10];15(3):382–92. Available from: http://www.pubmedcentral.nih.gov/articlerender.fcgi?artid=4059512&tool=pmcentrez&rendertype=abstract

62. HMP. Structure, function and diversity of the healthy human microbiome. Nature [Internet]. 2012 Jun 14 [cited 2014 Jan 20];486(7402):207–14. Available from: http://www.pubmedcentral.nih.gov/articlerender.fcgi?artid=3564958&tool=pmcentrez&rendertype=abstract

63. Schloissnig S, Arumugam M, Sunagawa S, Mitreva M, Tap J, Zhu A, et al. Genomic variation landscape of the human gut microbiome. Nature [Internet]. Nature Publishing Group, a division of Macmillan Publishers Limited. All Rights Reserved.; 2013 Jan 3 [cited 2014 Jan 20];493(7430):45–50. Available from: http://dx.doi.org/10.1038/nature11711

64. Markowitz VM, Mavromatis K, Ivanova NN, Chen I-MA, Chu K, Kyrpides NC. IMG ER: a system for microbial genome annotation expert review and curation. Bioinformatics [Internet]. 2009 Sep 1 [cited 2015 Feb 27];25(17):2271–8. Available from: http://www.ncbi.nlm.nih.gov/pubmed/19561336

65. Angly FE, Willner D, Rohwer F, Hugenholtz P, Tyson GW. Grinder: a versatile amplicon and shotgun sequence simulator. Nucleic Acids Res [Internet]. 2012 Jul [cited 2014 Nov 3];40(12):e94. Available from: http://www.pubmedcentral.nih.gov/articlerender.fcgi?artid=3384353&tool=pmcentrez&rendertype=abstract

66. Quail MA, Smith M, Coupland P, Otto TD, Harris SR, Connor TR, et al. A tale of three next generation sequencing platforms: comparison of Ion Torrent, Pacific Biosciences and Illumina MiSeq sequencers. BMC Genomics [Internet]. 2012 Jan [cited 2014 Jul 9];13:341. Available from: http://www.pubmedcentral.nih.gov/articlerender.fcgi?artid=3431227&tool=pmcentrez&rendertype=abstract

67. Rice P, Longden I, Bleasby A. EMBOSS: The European Molecular Biology Open Software Suite. Trends Genet [Internet]. 2000 Jun [cited 2014 Jan 28];16(6):276–7. Available from: http://www.sciencedirect.com/science/article/pii/S0168952500020242

68. Storey JD, Tibshirani R. Statistical significance for genomewide studies. Proc Natl Acad Sci U S A [Internet]. 2003 Aug 5 [cited 2014 Aug 17];100(16):9440–5. Available from: http://www.pnas.org/content/100/16/9440.short

